# Intraspecies warfare restricts strain coexistence in human skin microbiomes

**DOI:** 10.1101/2024.05.07.592803

**Authors:** Christopher P. Mancuso, Jacob S. Baker, Evan Qu, A. Delphine Tripp, Ishaq O. Balogun, Tami D. Lieberman

## Abstract

Determining why only a fraction of encountered or applied strains engraft in a given person’s microbiome is crucial for understanding and engineering these communities. Previous work has established that metabolic competition can restrict colonization success *in vivo*, but the relevance of bacterial warfare in preventing commensal engraftment has been less explored. Here, we demonstrate that intraspecies warfare presents a significant barrier to strain coexistence in the human skin microbiome by profiling 14,884 pairwise interactions between *Staphylococcus epidermidis* isolates cultured from eighteen people from six families. We find that intraspecies antagonisms are abundant, mechanistically diverse, independent of strain relatedness, and consistent with rapid evolution via horizontal gene transfer. Critically, these antagonisms are significantly depleted among strains residing on the same person relative to random assemblages, indicating a significant *in vivo* role. Together, our results emphasize that accounting for intraspecies warfare may be essential to the design of long-lasting probiotic therapeutics.

## Main

It is now clear that new strains are acquired throughout life in the gut^1–3^, oral^3^, vaginal^4^, and skin^5,6^ microbiomes. Despite this potential for transmission, the microbiomes of cohabitating adults do not homogenize^7^, with each individual harboring person-specific strains^2,8^. Neutral priority effects^9,10^, such as competition with established strains for metabolic^11–15^ or spatial niches^16,17^, have been explored as potential drivers of person-specificity. Alternatively, selection for specific species or strains could be driven by the immune system^18,19^, host behavior (e.g. diet^20^ or hygiene^21^), or other members of their microbiome^22^.

Intraspecies warfare is a compelling source of person-specific selection and barriers to strain colonization. Bacteria often actively antagonize other bacteria— from the same^23–25^ or different^26–29^ species— through a variety of contact-dependent and contact-independent mechanisms^22,30^. Both antimicrobial production and resistance can vary within species, often changing quickly through horizontal-gene-transfer^25,31,32^. To date, most work on interbacterial warfare in microbiomes has focused on antagonisms between different species^24,27,33^ and its potential to provide colonization resistance against pathogens^28,34^. However, given that niche overlap is highest within a species^35^, the selective advantage for inhibiting competing microbes is expected to be strongest when the competitor is from the same species^33,22,30^.

Here, we leverage *Staphylococcus epidermidis* in the facial skin microbiome as an ideal system for studying barriers to strain coexistence. Each person’s facial skin microbiome is typically colonized by several strains of *S. epidermidis* arising from distinct colonizing genotypes^6,17^; these compositions are dynamic, with each lineage persisting for an average of two years^5,6^. However, cohabitating individuals retain unique compositions despite this capacity for turnover^6^. Physical barriers to transmission are expected to be low for surface-dwelling microbes on skin, (as opposed to nasal or pore-dwelling microbes) necessitating an explanation for the person-specificity of these intraspecies communities.

*S. epidermidis* is known to produce a variety of interspecies antimicrobials^29,36–39^, and the accessory genome harbors an enrichment of antagonism-related biosynthetic gene clusters (BGCs) and defense genes (Supplementary Figure 1; Methods). Studies in the nasal microbiome show that people colonized by antimicrobial-producers can be protected from the pathogen *Staphylococcus aureus*^40,41,39^. We asked whether prevalent antimicrobial production uniquely targets *S. aureus*: could warfare be similarly common among commensal strains of the same species? If so, could this contribute to the yet unexplained person-specificity of commensal strain composition?

To test if intraspecies warfare is a major barrier to *S. epidermidis* strain coexistence, we examined antagonistic interactions among a collection of *S. epidermidis* isolates collected from extensively-sampled nuclear family members^6^. Ecologically-relevant warfare should result in fewer antagonisms between strains on a person than expected by random assembly of available strains (Figure 1A).

**Figure 1:**
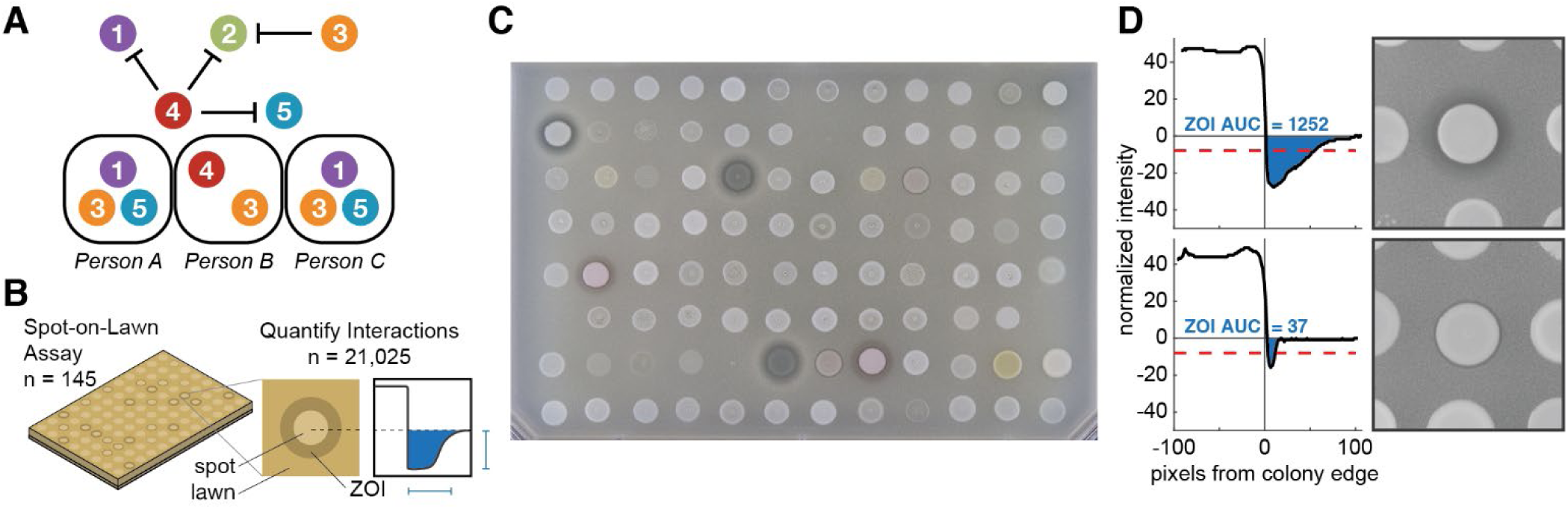
High-throughput assay of interbacterial warfare from skin microbiomes of related people. (A) Antagonism between different commensal strains of the same species (circles) could theoretically prevent transmission of certain strains despite frequent contact between individuals. Ecologically-relevant warfare would result in a depletion of antagonisms on individual people relative to random assortment. (B) We built a library of 122 *Staphylococcus epidermidis* isolates representing the strain diversity on healthy individuals from six families, and included 23 isolates from other skin species from these people. We screened every pairwise interaction (14,884 *S. epidermidis* intraspecies interactions plus 6,141 including other species) using high-throughput “spot-on-lawn” assays, in which producer cells are spotted on top of a lawn of receiver cells; we identified hundreds of cases of antagonism, which appear as Zones-Of-Inhibition (ZOIs; Methods). (C) Representative photograph of spot-on-lawn assay. Several *S. epidermidis* spots produce antimicrobials that antagonize the *S. epidermidis* receiver lawn, resulting in ZOIs of a range of intensities. Some spots on the plate are blank or were excluded for poor growth (see Supplemental Table 1 for array information). (D) We developed a computer-aided image analysis pipeline to identify even subtle ZOIs that pass the inhibition threshold (red dash) and quantify the strength of antagonism, determined by the Area Under the Curve of the intensity traces (AUC, blue; Methods).

Leveraging a collection of 1,870 sequenced *S. epidermidis* isolates from 18 people from 6 families, from which we also have metagenomic data^6^, we built a library of 122 isolates that capture genomic diversity on these individuals (Supplementary Table 1). This library includes 59 lineages, clusters of isolates separated less than 90 point mutations across their entire genomes^6^. Of these, 30 lineages were shared between family members and 29 lineages were unique to an individual. We also included 23 isolates of other skin bacterial species incidentally isolated from the same subjects. This library reflects ∼96% of the *S. epidermidis* relative abundance on these people as previously reported^6^ (Supplementary Figure 2, Supplementary Table 3), enabling efficient estimation of antagonism frequencies.

We conducted high-throughput solid-phase competition assays (“spot-on-lawn”^26^) to assess toxin production and sensitivity for each of 21,025 pairwise combinations of isolates (Figure 1B). For each isolate, we made a bait lawn by embedding cells into soft agar. We then grew the full library to mid-log phase in liquid media, spotted aliquots on top of bait lawns at high cell density, and allowed cells to grow until the bait lawns were opaque (Methods). A zone of inhibition (ZOI) was observed around ∼8% of spots, indicating the bait’s sensitivity to an antimicrobial produced by the spot (Figure 1C). We quantified the presence and size of ZOIs computationally (specifically the Area Under the Curve for intensity), enabling detection of even weak inhibition (Figure 1D).

## Antimicrobials are diverse and fast-evolving

A large percentage of *S. epidermidis* isolates antagonize at least one lineage from the same species (44%) or different species (70%) (Supplementary Figure 3). There is a wide distribution in the number of lineages antagonized; four lineages from different families each antagonize over half of all tested lineages. The percent of lineages inhibited by each antagonizer follows a roughly exponential distribution (Supplementary Figure 4) and we therefore we do not exclude any lineages from our analysis.

Both antimicrobial production and sensitivity are generally conserved at the lineage level but display no longer-lasting phylogenetic structure. Only 3.1% and 3.4% of possible interactions showed intra-lineage variation in antagonism or sensitivity, respectively (Supplementary Figure 3; Methods), and most of these remaining cases represent technical noise near the limit of detection (Supplementary Figure 5). The rarity of intra-lineage variation suggests minimal evolution of antimicrobial production or resistance over short on-person timescales. However, at broader within-species levels, antagonism is broadly distributed across the phylogeny with no apparent clustering (Figure 2A) and little correlation with sequence similarity (linear correlation R^2^ =0.27, *P*=0.046; Supplementary Figure 6). The lack of phylogenetic clustering suggests warfare profile changes on the order of decades, as lineages have a median distance of 416 point mutations from the nearest lineage (range 75-14,000). These rapid changes are consistent with either acquisition of antimicrobial production genes through horizontal gene transfer or frequent activation and deactivation of production through regulatory mutations.

**Figure 2:**
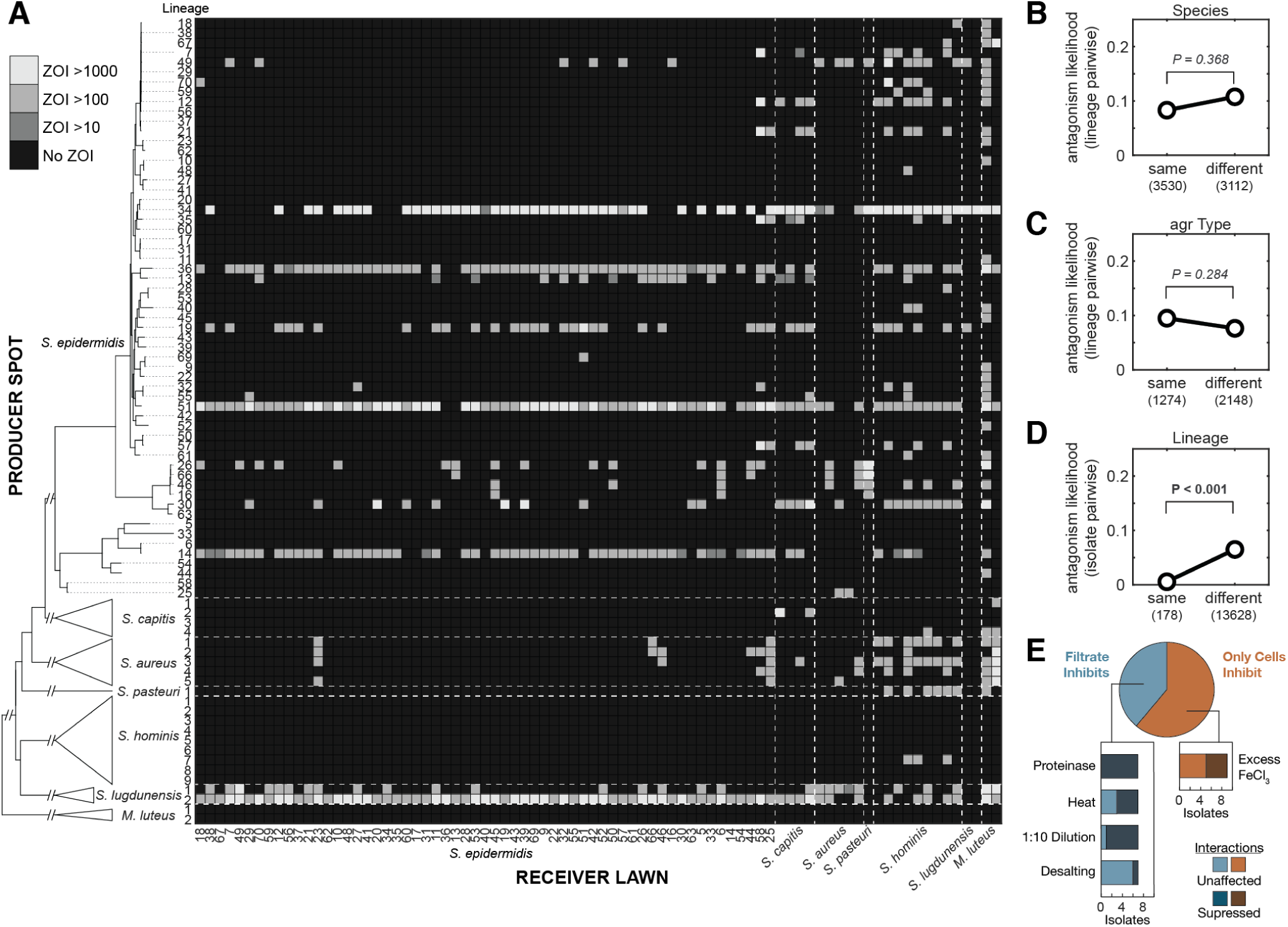
Intra-species antagonism is common and does not segregate at higher phylogenetic levels. (A) No evolutionary structure is observed in antimicrobial production (rows) or sensitivity (columns) beyond the lineage level, indicating that antimicrobial production and resistance are fast evolving and not conserved. The heatmap depicts the AUC for each ZOI calculated for pairwise interactions between skin isolates dereplicated to the lineage level (Methods). Each pair is screened twice, with each isolate serving as both spot and lawn. Inhibition of a receiver lawn by a producer spot is indicated by a lighter square, with intensity indicating the size of the ZOI. Lineages and species are sorted by phylogenetic similarity (see tree at left, not to scale beyond *S. epidermidis*), and dashed lines on the heatmap species boundaries. Supplementary Figure 3 depicts this data without dereplicating *S. epidermidis* isolates into lineages. (B-C) The likelihood of observing antagonism between two lineages does not significantly differ between members of the same or different (B) species or (C) quorum sensing variant (agr type). Significance p-values are derived from two-sided permutation tests with shuffled group labels (Methods). (D) Conversely, intra-lineage antagonism is negligible compared to inter-lineage antagonism. (E) To assess the mechanisms of antagonistic interactions, we repeated the screen using cells and filtrates from 18 *S. epidermidis* producers against an array of bait lawns chosen based on interaction profiles (Supplementary Figure 7). Filtrates were subjected to a variety of pre-treatments. Nine interactions that did not inhibit in the filtrate were retested on media containing excess iron to identify iron-suppressible interactions. The results suggest a large variety of interaction mechanisms which are not neatly predictable from genomic context in this screen (Supplementary Figure 8).

We observe no significant difference between frequencies of interspecies and intraspecies antagonism when accounting for structure in the interaction matrix (*P* = 0.368, permutation test; Figure 2B; Methods); moreover, some antagonists participate in both intra– and inter-species killing. There was also no difference in antagonism whether lineages used the same or different quorum sensing variant (agr type, *P* = 0.284, permutation test, Figure 2C), suggesting that intraspecies communication is not a major determinant of sensitivity. We observe a negligible number of intra-lineage antagonisms (*P* < 0.001, permutation test; Figure 2D), consistent with the rarity of intra-lineage variation. These observations indicate that it may be impossible to predict production or sensitivity to intraspecies antimicrobials using higher-order phylogenetic signals or single marker genes.

The large number of antagonism patterns observed suggests a diverse pool of antimicrobial molecules (at least 34 different antagonism patterns, Supplementary Figure 7) with no universally prevalent molecules in *S. epidermidis*. For 18 isolates chosen based on the abundance and diversity of interaction patterns (Supplementary Figure 7; Methods), we repeated screening using filtered culture supernatants that were either treated with heat, proteinase-K, removal of small molecules by desalting, dilution, or not treated (Figure 2E, Supplementary Figure 8A, and Supplementary Table 4). Seven of these isolates had activity in the supernatant; all of these antagonisms were proteinase sensitive and are therefore likely mediated by peptides or proteins. The radii of the cell-dependent ZOIs suggest diffusible molecules lacking efficacy in the supernatant for reasons of stability, inducibility, or concentration rather than contact-dependence. We repeated spot-testing to test for iron-suppression^29^, which modified four out of the nine cell-dependent interactions tested (Supplementary Figure 8B; Methods). Together, these results suggested a wide variety of competitive mechanisms, including peptides, an iron-chelator^42^, and other proteins (Fig. 2E). Importantly, the most antagonistic lineages had diverse mechanisms of interaction (Supplementary Tables 4 and 5). Multiple lineages showed patterns indicative of multiple effector types (Supplementary Figure 8C).

Some antagonist lineage genomes contain potential biosynthetic clusters (BGCs) for known products (including gallidermin^37^ and epicidin^38^) and unclassified products (Supplementary Table 5), supporting a role for horizontal gene transfer in driving rapid within-species changes. However, not all isolates with the same BGC had the same interactions, complicating the systematic linking of interactions and BGCs. While some of these cases are artifacts of fluctuation around our detection threshold, many other cases are robust and therefore implicate the presence of multiple antimicrobial molecules per isolate, the incomplete expression of all BGCs, or both. Moreover, not every antagonist had an identified BGC, highlighting the likelihood that new classes of antimicrobials remain to be discovered.

Incomplete or variable BGC expression could be driven by environment-dependent regulatory pathways. We therefore repeated the screen, initially performed Tryptic Soy Agar (TSA) with a media designed to more closely resemble skin: minimal M9 media supplemented with artificial sweat^14,43^ (M9+S; Supplementary Figure 9). Overall, we observed that 91% of interactions remained the same in both media, though antagonistic interactions did vary between the two media. Thus, only 23% of antagonisms observed in TSA were also observed in M9+S, and 43% of antagonisms observed in M9+S were observed in TSA (Supplementary Figure 10A). M9+S supported significantly less growth than TSA; as such, some of these observed differences may reflect decreased assay sensitivity in low-growth conditions, rather than true changes in expression profiles. Moreover, recent studies have shown that antimicrobial production genes are expressed *in vivo*^43,44^, raising the likelihood that these interactions are ecologically relevant.

## On-person depletion of warfare

To test for ecological relevance of these antimicrobial interactions, we looked to see if antagonistic interactions were depleted among lineages present on a same person at a single timepoint, relative to randomly chosen lineages. We assessed the Antagonism Frequency (AF) for each sample, a metric that accounts for lineage abundances (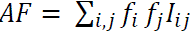 where *f* represents the relative abundance of that lineage in that sample and 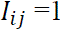 if lineage i antagonizes lineage j; see Methods). Mean AF computed on the average composition across all samples in our collection is 0.14; this can be roughly interpreted as the likelihood of antagonism between a pair of randomly chosen bacteria out of the *S. epidermidis* population. Despite this expectation, we find that the majority of samples have AF values of zero, suggesting a depletion of on-person antagonism (Supplementary Figure 11A). Antagonisms remain depleted when each subject is counted just once, using mean AF across samples (Figure 3A).

**Figure 3:**
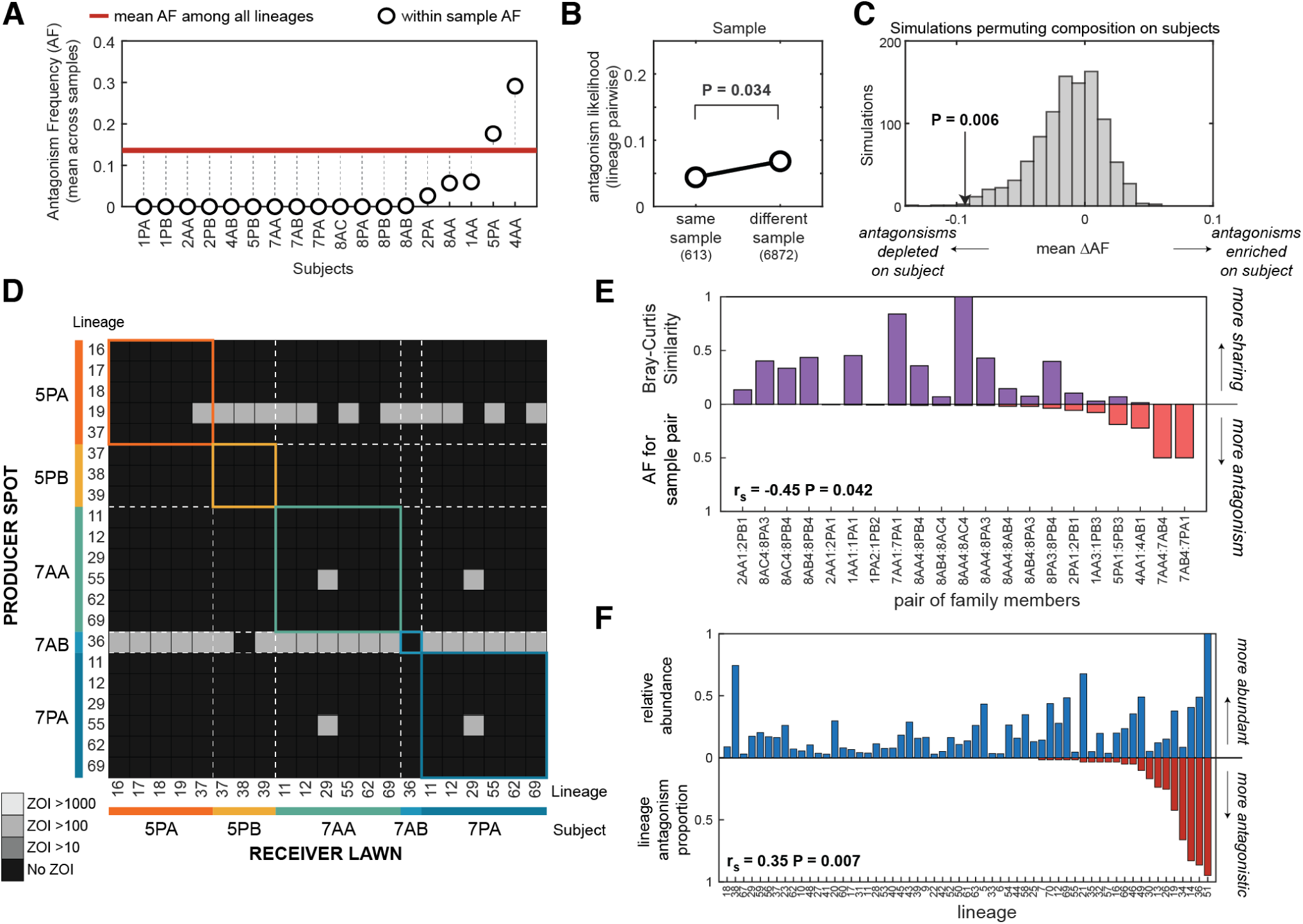
Antagonism is depleted among lineages co-residing on individuals. We combined the results of the antagonism screen with isolate-inferred relative abundances of each lineage on each person (Supplementary Figure 2 and Supplementary Table 3) to study the ecological relevance of these interactions. (A) The abundance-weighted Antagonism Frequency (AF) of interactions between co-resident lineages in samples (e.g. same person, same timepoint) is depleted relative to AF calculated across all subjects pooled together (red line). Each datapoint depicts the mean AF across samples from a subject. Subject codes indicate family and subject (e.g. Subject 1PA is from family 1). (B) Lineage pairs that occur in the same sample are less likely to antagonize each other than lineages from different samples, relative to permutation tests where sample origin is shuffled; this analysis maintains antagonism structure (e.g. wide distribution of percent of isolates antagonized). (C) Simulations of a null model that incorporate lineage composition on subjects (e.g. uneven relative abundances of lineages) as well as antagonism structure also support a significant depletion of antagonism in the observed AF (arrow) of coexisting lineages on individuals. In each simulation, the antagonism matrix is kept constant but lineage identities are shuffled across all subjects in our study. As each simulation has a different expected AF, we can compare the mean difference between each subject’s AF (i.e. white datapoints from A) and the expected AF (i.e. red line from A) and the mean difference from each simulation (mean ΔAF). (D) A section of the lineage-level heatmap from Figure 2A, sorted to show only lineages present on subjects in two families (see Supplementary Figure 12 for all families), highlights that lineages on a person often antagonize lineages on other family members. Intensity indicates AUC for the ZOI, and lineages are repeated if present on multiple subjects. (E) The Bray-Curtis Similarity between the lineage composition of sample pairs (purple) is negatively correlated with the AF calculated for each sample pair after dereplication (red; Spearman’s rank correlation), supporting a role for antagonism in preventing transmission. (F) Lineages that antagonize a higher proportion of other lineages (red) reach higher abundance on individuals (blue; Spearman’s rank correlation). Each lineage’s antagonism proportion was calculated by counting the other lineages it produced a ZOI on and dividing by the total number of lineages.

We compared the likelihood of antagonism between lineages that co-occur in the same sample versus lineages that never co-occur, revealing a statistically significant depletion of antagonism between co-resident lineages (*P* = 0.034, permutation test; Figure 3B). However, this analysis does not account for variation in lineage composition across individuals, which ranged from a single highly dominant strain to nine coexisting strains (Supplementary Figure 2). We therefore applied permutation tests in which lineage labels are shuffled across all samples in the population in order to simulate different compositions, which confirmed a significant depletion of antagonism on individuals (*P* = 0.006, permutation test; Methods; Figure 3C). The depletion of on-person antagonism is robust to whether interactions were scored on TSA, M9+S, or in any combination of these screens (Supplementary Figure 10B-E). The depletion of on-person interactions was reproducible across a variety of permutation tests with varying assumptions (described in Methods), including elimination of sharing of lineages across families or shuffling each subject independently from their family members (*P* < 0.032, permutation tests; Supplementary Figure 11C-F). The robustness of this result to varying media conditions, scope of transmission, and person-specificity support the *in vivo* relevance of these interactions.

The ecological signature of antagonism is easily observable when the interaction network is sorted by subject (Figure 3D, Supplementary Figure 12) or when lineage compositions are colored by the antagonism ability of each lineage (Supplementary Figure 13). Subjects carrying antagonist lineages have substantially different *S. epidermidis* compositions than their family members, who carry sensitive lineages (e.g. 5PA, 7AB; Figure 3D). Accordingly, subjects colonized by lineages antagonistic to lineages of family members share fewer lineages with those family members (r_s_ = –0.45, *P* = 0.042, Spearman Rank Correlation; Figure 3E), suggesting that antagonism poses a barrier to colonization by incoming strains. Notably, other assessments of *S. epidermidis* diversity, including quorum-sensing variants^45^ or phylogenetic groupings (Supplementary Figure 14), do not display these person-specific ecological signatures.

This indication of the importance of within-species warfare suggests that antagonistic lineages have increased fitness on human skin relative to non-producing strains. Accordingly, lineages that antagonize many others generally reach higher relative abundances of the *S. epidermidis* community on individual people (r_s_ = 0.35, *P* = 0.007 Spearman Rank Correlation; Figure 3F). This correlation between antagonism and *in vivo* abundance is stronger than that between *in vitro* growth rate and *in vivo* abundance (r_s_ = 0.26, *P* = 0.045, Spearman Rank Correlation; Supplementary Figure 15C). Since antagonism provides a selective advantage on individuals, but this does not translate to high global prevalence of any specific antagonist or antimicrobial gene cluster, there must either be tradeoffs in overcoming other colonization barriers (e.g. priority effects) or frequent evolution of resistance undetected in our sampling. We did not find evidence of strong tradeoffs for antimicrobial production *in vitro*, with producers and non-producers having comparable growth rates (*P* = 0.190, Spearman Rank Correlation; Supplementary Figure 15D); however, this does not rule out the possibility of tradeoffs for production *in vivo*. In addition, it is probable that nutrient competition^14,46^, immune evasion^47^, or other selective forces could enable resistant strains to outcompete antimicrobial producers^24,48^.

## Rare on-person evolution of sensitivity

While 97% of interactions did not exhibit intra-lineage variation, a few outlier lineages exhibited large differences in antagonism sensitivity profiles (Methods), suggesting on-person evolution of resistance or sensitivity. Surprisingly, the phylogenetic tree structures were indicative of on-person evolution of sensitivity from a resistant ancestor (Supplementary Figure 16). As these “idiosyncratic” isolates formed a minority of their lineages, these interactions were excluded from other analyses presented in this study, though our results are robust to their inclusion (Supplementary Figure 16A).

Interestingly, two “idiosyncratic” isolates with derived sensitivity (isolate 3 from lineage 20, 20.3 found on subject 2PA; and isolate 3 from lineage 37, 37.3, found on subject 5PA) were vulnerable to the same group of antagonists (Figure 4A). We therefore asked whether they were caused by similar mutations, which would be evidence for parallel evolution. Despite fewer than 90 mutations separating isolates in each lineage, we found a clear commonality: a nonsense mutation in *vraG* in isolate 20.3 and a frameshift in *vraF* in isolate 37.3 (Supplementary Figure 16). Deletions in the *vraFG* signal-transduction complex^49^ or its downstream pathways^50,51^ are known to increase sensitivity to cationic antimicrobials produced by bacteria and humans; these mutations suppress modifications to wall-teichoic acid^51^ and phosphatidylglycerol^50^ that increase cell envelope charge. Consistent with this expectation, idiosyncratic isolates were more sensitive to the cationic antimicrobials we tested (Figure 4D), but not mildly charged ones (Supplementary Figure 17). Idiosyncratic isolates also had increased sensitivity to lysostaphin, an enzyme targeting the cell wall (Figure 4D).

**Figure 4:**
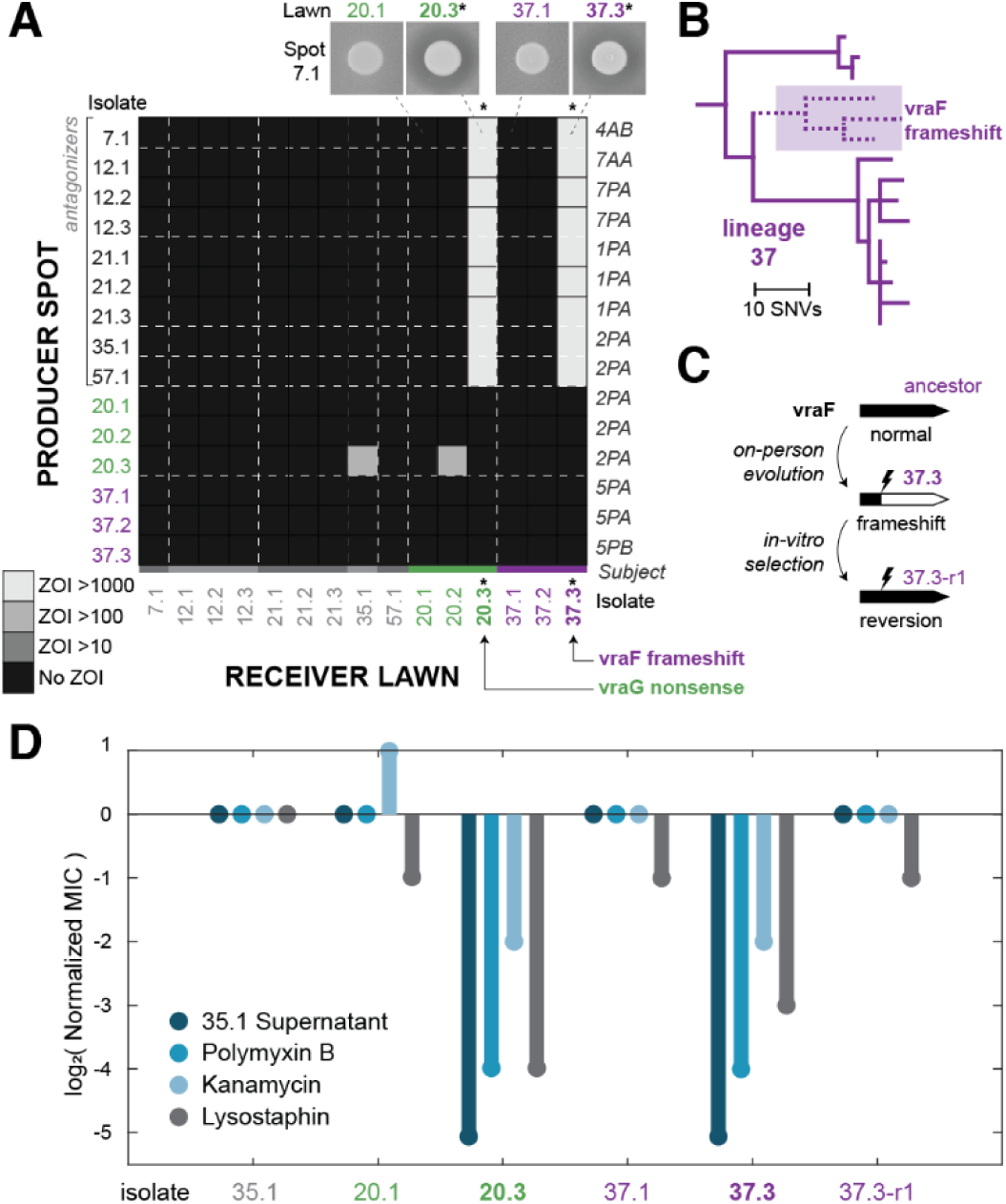
On-person evolution drives sensitivity to multiple antimicrobials. (A) Two lineages show significant intra-lineage variation, each containing isolates with sensitivity to the same producers, depicted using a section of the isolate-level heatmap (from Supplemental Figure 3). Isolates from the same lineage are indicated by decimals and color. Intensity indicates AUC for the ZOI. The two idiosyncratic isolates depicted (denoted by *) have different sensitivity profiles from the rest of their lineages and also have loss-of-function mutations in the vraFG pathway, as labeled. (B) The phylogeny of isolates in lineage 37 indicates that the vraF frameshift and associated sensitivity was acquired, not ancestral. Lineage 20 has a similar pattern of recently acquired sensitivity (see Supplemental Figure 16). (C) By selecting on polymyxin B, we cultivated revertant isolate 37.3-r1 which corrected the vraF frameshift and has an ancestral resistance profile. Genome sequencing confirmed this reversion (Methods). (D) Isolates with vraFG mutations, but not their ancestors or the revertant, are sensitive to cationic antimicrobials, lysostaphin, and filtered supernatant from isolate 35.1. Minimum Inhibitory Concentration (MIC) assay results were normalized to isolate 35.1 and raw values are available in Supplemental Figure 18.

To support the causative role of *vraFG* mutations driving sensitivity, we plated isolate 20.3 and 37.3 on high levels of polymyxin B to select for resistant colonies (Figure 4C; Methods). While we did not get resistant colonies from 20.3, we isolated ten resistant isolates derived from 37.3, two of which had corrected the *vraF* frameshift. Both revertants had an ancestral resistance profile (see isolate 37.3-r1 in Figure 4C-D and isolate 37.3-r2 in Supplementary Figure 17). These results confirm that the observed sensitivity was driven by *de novo* mutations acquired on individuals that break the *vraFG* pathway.

The parallelism between the two lineages strongly suggests that the loss of *vraFG* is advantageous on these hosts. Moreover, a second lineage (58) on one of the subjects with an idiosyncratic lineage (2PA) showed the same pattern of sensitivity (Supplementary Figure 7) and also had nonsynonymous mutations in the same genetic pathway (Supplementary Figure 16). Together, these results raise the possibility that *vraFG* loss creates a tradeoff between warfare sensitivity and some other, advantageous, phenotype. The *vraF* mutant 37.3 grows slower *in vitro* than its full-length *vraF* revertant 37.3-r1 (Supplementary Figure 15B) suggesting growth advantages are insufficient to explain this selection, but a growth tradeoff may still be present *in vivo*. Cell envelope modifications often affect phage susceptibility, including multiple wall teichoic acid modifications recently characterized in *Staphylococci*^52,53^; mutant 37.3 displayed partial resistance to a phage which supports the plausibility of this tradeoff (Supplementary Figure 18). However, we were unable to find this specific phage in metagenomic samples from the subjects carrying vraFG mutants (Methods), nor isolate other phages from these subjects (Methods). Thus, more work is needed to understand the drivers of on-person evolution of sensitivity, including the possible roles for nutrient competition^14^, immune evasion^47^, or other *in vivo* selective forces.

## Discussion

Here we show that interbacterial warfare plays a significant role in determining commensal survival on human skin. These observations reconcile the accessibility of the skin to new *S. epidermidis* colonizations^6^ with the person-specific nature of strain composition^5,54,45,6^.

Given the advantage that warfare seems to confer, why is there so much variation in mechanism and the amount of production and sensitivity across the population? While the potential reasons are numerous and complex, our results point to the existence of selective tradeoffs for antimicrobial resistance; such tradeoffs may also exist for production of different weapons. Antimicrobial exporters and cell envelope components could be targeted by phage^53,55^ and the immune system^19,47^, imposing harsh selective pressures against antagonists and resistors on certain hosts. While we find no consistent growth deficit for production or resistance here, this has been reported for other antimicrobials^46^. Furthermore, fitness deficits induced by the cost of production may also be more pronounced *in vivo* and lead to rock-paper-scissor dynamics among producer, resistant, and sensitive populations as have been shown in simulations and *in vitro*^24,48^. Lastly, priority effects^9,10^ and selection on orthogonal phenotypes^14,15^ may play important roles.

This evidence of *in vivo* exclusion by antagonists suggests strain mixing at length scales over which antimicrobials can act. Unlike *C. acnes,* which is thought to primarily grow in pores with low lineage diversity^17^, *S. epidermidis* is generally thought to live on the surface of skin^56^ and mix with other strains and species. However, the fine-scaled spatial structure of *S. epidermidis* on skin remains to be explored. Determining whether strains are evenly mixed or only compete at the borders of small, overlapping clonal patches will require the application of additional sampling^57^ or imaging methods^58^.

While other studies have identified trends between phylogenetic distance and probability of antagonism^24,27,33,59^, we find little signal here. Instead, our observations are consistent with rapid horizontal gene transfer (over the last couple decades) and molecules that mediate both interspecies and intraspecies warfare; these phenomena both make it difficult to interpret the relationship between phylogenetic distance and antagonism. Whether intraspecies or interspecies^29,36,28,60,61^ antagonism provides the primary selective advantage for production is unclear. Given the high metabolic overlap among strains of a species^33,35^, it is possible that interspecies killing is simply a byproduct of intraspecies killing. Experiments enabling higher throughput, combined with deep strain profiling from many species on the same set of people, will be required to address the relative roles of intraspecies and interspecies warfare on humans. Regardless our results show that intraspecies warfare contributes to strain composition *in vivo*.

The high turnover of antimicrobial production and sensitivity highlight that whole-genome resolution may be required to predict the ability of a person’s microbiome to resist pathogens or probiotics^2^. However, even given perfect genomic information, understanding the genetic basis of production and resistance will be critical. Particularly given the potential for tradeoffs between resistance to the host^62,63^, phage^52^, and antimicrobials, the specific mechanisms of resistance will be paramount for predictive modeling. The diversity of antagonistic patterns observed here precludes straightforward association of interactions to genetic elements, highlighting the need for novel approaches to identify and characterize new antimicrobials, BGCs, and resistance determinants^64–67^.

Many studies have focused on overcoming metabolic^68^ and immune^47^ barriers to engraftment in human microbiomes. Our results assert that interbacterial antagonism should be considered alongside these forces when designing long-lasting microbiome-targeted therapies. Longitudinal or interventional colonization studies are needed to provide more direct evidence to understand if warfare acts at the level of transmission or persistence, and whether it drives the acquisition of resistance *in vivo*. The depletion of antagonisms at single timepoints seen here suggests that long-term probiotic success^2^ will be enhanced by eliminating pre-existing microbes, confirming resistance to resident toxin-producers, or bet-hedging with multiple probiotic formulations. Moreover, after overcoming these barriers, engrafted probiotics that actively protect their niches could ensure long-lasting therapeutic benefits.

## Methods

### Collection and classification of genomically diverse S. epidermidis

Isolates were selected from the collection of *S. epidermidis* created in Baker et al^6^; methods and data relevant to the study cohort, isolate collection, sequencing, metagenomic analysis with PHLAME^69^, and transmission dynamics can be found in that study. In short, 2,025 *S. epidermidis* isolates were clustered into 78 lineages based on core genome single nucleotide polymorphisms; lineages had a median distance of 416 core genome point mutations from the nearest lineage (range 75-14,000)^6^. Isolates within a lineage have fewer than 90 point mutations across their core and lineage-shared accessory genome^6^. The core genome mutational clock rate was measured to be 4.37 mutations/genome/year^6^, indicating that lineages share a recent common ancestor no more than ∼10 years ago. PHLAME^69^ was used to confirm that lineage abundances estimated by culturing did not have significant culture bias^6^; we therefore used isolate-based lineage abundances because deep metagenomic coverage of *S. epidermidis* was not available for every sample.

In this work, we focused on 6 families chosen because they each had >100 isolates of *S. epidermidis*, and included samples with >20 isolates per sample. In total, 122 representative isolates were selected from each lineage present on subjects from these families. When possible, multiple isolates were chosen from a lineage in order to have isolates from multiple subjects and with different mobile element content. Mobile element content was defined using mapping to lineage co-assemblies as previously described^6,17^. In addition, 23 isolates from other Staphylococcus species and *Micrococcus luteus* from these families were also included. Isolate details are found in Supplementary Table 1. Rothia and Streptococcus isolates were also originally included in these arrays, but were not analyzed due to poor growth in the conditions of the assays. These were later replaced by 11 new *S. epidermidis* isolates, see Supplementary Table 2.

### Sequencing and genome assembly

All *S. epidermidis* isolates were sequenced as part of Baker et al^6^. Additional whole-genome sequencing was performed on the 23 non-epidermidis isolates included in this study. DNA extraction was performed using the PureLink Genomic DNA kit (Invitrogen), substituting 40µg/mL lysostaphin (prepared in PBS) and ReadyLyse (Biosearch Technologies) for the lysozyme step in the manufacturer’s protocol. Shotgun sequencing libraries were generated following the HackFlex protocol^70^. 150 bp paired end sequencing reads were generated on an Element Aviti system. Sequencing reads were assembled using SPAdes v3.15.3^71^ and annotated using Bakta v1.8.1^72^. The Snakemake^73^ pipeline used for genome assembly and annotation is available on GitHub (see Code Availability).

### Culture conditions

Unless otherwise noted, all isolates were grown in Tryptic Soy Broth (TSB) (Neogen NCM0004A) aerobically at 37°C, with shaking at 250 RPM if in liquid culture, or on Tryptic Soy Agar (TSA) with 1.5% agar at 37°C without shaking if on solid culture. For all experiments, mid-late exponential phase cells were prepared by diluting 5 µL of overnight culture into 495 µL of fresh media and growing these subcultures for 4-6 hours.

### Spot-on-lawn assay in TSA

Glycerol stocks of single isolates were thawed, then 5 µL of glycerol stock was diluted into 495 µL of fresh TSB + 20% glycerol, arrayed into two 96-well plate arrays (Supplementary Table 1), and aliquoted into 10 µL replicate plates. For each experiment, a replicate plate was thawed; thawed plates were never reused. Overnights were prepared by diluting 5 µL of glycerol aliquot into 495 µL TSB in 2 mL deep-well blocks. Mid-late exponential phase cells were prepared by diluting 5 µL of overnight culture into 495 µL of fresh media and growing these subcultures for 4-6 hours.

For lawn preparation, soft Tryptic Soy Agar with 1% agar was autoclaved for 20 mins, then allowed to cool to 55°C. TSA was then further cooled to 42°C in a water bath (remining molten). For each lawn isolate, optical density of mid exponential phase cells was measured. Roughly 2.5 x 10^7^ cells (adjusted to 0.5 x 10^7^ for *S. aureus* to offset higher growth of this species) were added to 50 mL conical tubes along with 35 mL of molten soft TSA. Tubes were mixed by inverting 4 times, then poured into Nunc OmniTrays (Thermo Scientific 242811). Plates were allowed to cool for up to 2 h. While plates cooled, spot cultures were prepared by normalizing the OD of late log-phase sub-cultures to OD 0.1 (roughly 6 x 10^7^ CFU/mL) in fresh TSB. A 96-well pipettor (Avidien MicroPro300) was used to dispense 2 µL of spot culture onto each plate. After spots dried (∼15 mins), plates were bagged to prevent evaporation and incubated at 37°C for 24 h then photographed. To more clearly resolve ZOIs by increasing the turbidity of lawn cultures, plates were allowed to grow stored at 21°C for 48 h (to avoid desiccation in the incubator), and photographed again.

### Imaging and image analysis

Photographs were taken after 24 h and 72 h of growth. Plates were placed into a custom darkbox with diffuse lighting from the sides and the camera suspended above the plate. Photographs were taken at 6000 x 4000 pixel resolution on a Canon EOS Rebel SL2. Manual focus was determined a single time each day. All images with metadata have been uploaded to figshare (see Data Availability).

Image analysis was performed using a custom MATLAB script applied to greyscale images. Spots, their centers, and their edges were identified as regions of interest (ROIs) centered at 96 points using the imfindcircles function, with any questionable calls flagged for manual ROI definition. The median intensity of pixels for each radial distance from the identified edge was calculated for each spot. These values were corrected for local illumination using the intensity of regions between spots, defined as >180 pixels from any spot center. Spot centers, radii, intensities and variance were stored to allow the user to choose different thresholds to define ZOIs and are available on GitHub (see Code Availability). For the TSA results presented here, thresholds were set to an intensity depth of 8 (arbitrary intensity units) and Area Under the Curve (AUC) of 50 (arbitrary intensity units x pixels). Any spots exceeding either of these thresholds were classified as positive. These thresholds were chosen to be conservative and more prone to false negatives than false positives; among replicate spots of the same isolate pairs, ZOIs called in one spot were called only 93% of the time in replicate spots, with more errors near the limit of detection. The major results are robust to alternative threshold choices (Supplementary Figure 5). The dereplicated isolate-level interaction matrix from the TSA screen is available in Supplementary Table 6.

When dereplicating isolates into lineages, the maximum ZOI observed between a pair of lineages was chosen as representative. This choice was made due to our conservative ZOI calling pipeline. One case of self-inhibition by lineage 51 was overwritten and rescored as non-inhibitory since it was not reproducible across replicates. Calls for plates with poor growth or for spots in areas of glare were masked from subsequent analyses (treated as no data). To calculate intra-lineage variation, we considered only lineage pairing with more than with more than one producer or receiver isolate (for antagonism or sensitivity, respectively). We then divided the number of lineage pairings where isolates disagreed in ZOI call (e.g. ZOI present for one isolate but not another in the same lineage) by the total number of interactions in these filtered datasets.

### Preparation of M9 + Sweat media

M9 + Sweat media was adapted from Swaney et al^14^, with a few changes. 8X artificial sweat was prepared exactly as described in Swaney et al and filter sterilized. Next, a 5X solution containing 56.4 g/L M9 salts, and 0.5 g/L glucose were autoclaved for 20 min. Next, these solutions were mixed to 1X working concentration and supplemented with 0.1 mM CaCl_2_, 2 mM MgSO_4_, 50 nM CoCl_2_, 100 mg/L l-arginine, 200 mg/L l-proline, 2 mg/L thiamine-HCl, 2 mg/L nicotinic acid, 2 mg/L calcium pantothenate, and 2 mg/L biotin. For liquid media, this final composition was prepared in sterile distilled water, whereas solid media was prepared using 1% autoclaved agar (10 g/L). Unlike Swaney et al^14^, instead of adding additional glucose at this stage, we added lactate to a final concentration of 10 mM, as lactate is a more common carbon source on skin.

### Spot-on-lawn assay in M9 + Sweat

The spot on lawn assay was repeated using the same general protocol as in TSA (see Spot-on-lawn assay in TSA), but using a different media composition in the following steps. Overnights were still prepared by diluting 5 µL of glycerol aliquot into 495 µL TSB in 2 mL deep-well blocks. Mid-late exponential phase cells were prepared by diluting 5 µL of overnight culture into 495 µL of M9 + Sweat media (instead of TSB) and growing these subcultures for 4-6 hours. Soft M9 + Sweat agar was prepared using 1% agar that had been cooled to 55°C. M9 + Sweat agar was then further cooled to 42°C in a water bath (remaining molten). Spot cultures were prepared by normalizing the OD of late log-phase sub-cultures to OD 0.1 in fresh M9 + Sweat media (instead of TSB). Image analysis was performed as before, but thresholds had to be changed to account for reduced growth: intensity depth threshold was changed to 4 (arbitrary intensity units) and AUC threshold was changed to 0. The dereplicated isolate-level interaction matrix in M9 is available in Supplementary Table 7 (also see Supplementary Figures 9,10).

### Statistical analysis of antagonism frequency

Weighted antagonism frequency (AF) was defined for each sample (Equation 1), where *f* represents the relative abundance of that lineage in that sample and 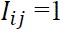 if lineage i antagonizes lineage j.

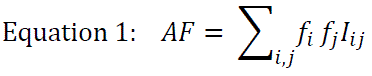

To define expected AF across all subjects in our cohort we averaged lineage abundances across all subjects to construct a population abundance vector. Weighting by abundance means that AF could be interpreted as the frequency of interaction between two cells on the same individual, as opposed to two lineages on the same individual. Since not all samples had sufficient sequencing depth for metagenomic analysis with PHLAME, we used relative abundances inferred from isolates, which have good agreement with metagenomic estimates^6^. The isolate-inferred relative abundances for each lineage are available in Supplementary Table 3.

We performed permutation tests by simulating scenarios that retain the structure of the interaction matrix and incorporate the uneven lineage distribution in each sample (Figure 3C; Supplementary Figure 11). For each simulation, we shuffled lineage labels without replacement in the relative abundance matrix (composed of one row for each sample with relative abundance f_n_ for every lineage in columns).

In order to compare across simulations when each has a different expected overall AF, Delta Antagonism Frequency (ΔAF) was defined as the difference between the per-sample AF and expected AF calculated from all lineages in our dataset. Mean ΔAF values were calculated for each subject by averaging across samples, and mean ΔAF values were calculated for each family by averaging across subjects. The p-value was defined as the fraction of simulations with less on-person antagonism (less negative ΔAF) than observed in our data.

By varying the method of shuffling, we simulated different transmission scenarios. First, we ran simulations in which the rows of the relative abundance matrix were shuffled once across all lineages in our collection. This simulation tests for depletion of antagonism while maintaining the same degree of lineage sharing seen between members of the same family (Figure 3C, Supplementary Figure 11C). Next, we ran simulations in which the rows of the composition matrix were shuffled, AF values were calculated for samples of the first subject, then rows were shuffled again before each new subject. This method broke the relatedness between individuals in the same family, while retaining similarity between samples from the same subject, thereby testing depletion of antagonism under the assumption that lineages are acquired from the environment independent of family (Supplementary Figure 11D). Third, we ran simulations that assume each family is only exposed to lineages found on their family members, shuffling the rows of the relative abundance submatrices composed only of lineages present on each family (Supplementary Figure 11E). Finally, we shuffled the family submatrices between each subject, which limits each subject to lineages from their family, but alters lineage sharing between family members (Supplementary Figure 11F). All of these simulations resulted in p-values < 0.05 (Supplementary Figure 11C-F).

### Mechanism screen for antagonistic interactions

Twenty isolates were picked for a mechanistic follow-up screen: eighteen isolates from the most antagonistic lineages (which each inhibited more than four other lineages) as well as two negative control isolates that lacked antimicrobial genes but were suspected to have active prophages (in order to distinguish antagonism from phage activity). Hierarchical clustering was performed using the MATLAB clustergram function (which uses Euclidean distance) in order to select 54 representative bait isolates (Supplementary Figure 7). Spot-on-lawn assays were repeated using this representative collection, with five additional supernatant variations for each antagonistic isolate. Supernatant from overnight cultures was passed through 0.22 µm filters (EMDMillipore MSGVS2210), and either (a) applied directly or treated with: (b) 3 h incubation with 19 µg/mL proteinase K (GoldBio P-480-SL2), (c) 30 min incubation at 95°C, (d) column-based desalting (Thermo Scientific 88512), or (e) 1:10 dilution into fresh TSB media. All supernatants were stored overnight at 4°C prior to spotting on bait lawns. Antagonisms were classified as cell-associated or filtrate-associated depending on whether ZOIs appeared around filtrate spots in addition to cell spots (Supplementary Figure 8).

To screen for iron-related mechanisms, a third round of spot-on-lawn assays were performed on 12 isolates: 9 with cell-associated interactions (no activity in the filtrate assays above), 2 with filtrate associated interactions and 1 *S. lugdunensis* isolate. Due to space constraints not every isolate from the mechanism screen could be tested, so we prioritized cell-associated isolates from different parts of the *S. epidermidis* phylogeny. Spot cultures and supernatants (untreated, heat, or proteinase treated) were prepared from cultures pre-grown in TSB or TSA + 90 µM FeCl_3_ and plated on either soft TSA or soft TSA + 90 µM FeCl_3_. Pre-growth condition did not alter inhibition phenotypes and filtrate-associated interactions had no altered phenotypes in any condition. Therefore, iron suppression was classified in cases where cell-associated ZOIs on the TSA plate were not observed on the TSA + 90 µM FeCl_3_ plate (Figure 2E, Supplementary Figure 8B, Supplemental Table 4).

### Identification of putative biosynthetic gene clusters

Published *S. epidermidis* lineage assemblies from Baker et al^6^ and the 23 non-epidermidis isolate assemblies from this work were annotated using Bakta v1.8.1 and screened against specialist databases using AMRFinder^74^, VFDB^75^, and Defense Finder^76^. Cluster of Orthologous Genes (COG terms) were assigned using eggnog-mapper v2^77^. Biosynthetic gene clusters (BGCs) were predicted for each lineage using antiSMASH^78^ version 7 with the version 7.0.0 database, and clustered using BiG-SCAPE2^79^ using the “--classify legacy” option. All BGCs analyzed are listed in (Supplementary Table 5). The Snakemake and local analysis scripts are available on GitHub (see Code Availability).

There were a few cases of intra-lineage differences in the presence or absence of putative antimicrobial BGCs (listed in Supplementary Table 5). However, isolates carrying the antimicrobial BGC did not clearly differ in antagonism profile from isolates that lacked those BGCs (e.g. 26.1 and 26.3 carrying lactococcin-like elements and 26.3 lacking lactococcin). These results highlight the difficulty in ascribing any particular antagonism to a particular genetic element without in-depth mechanistic studies.

### Mutational analysis of idiosyncratic isolates exhibiting intra-lineage variation

As a reminder, when dereplicating isolates into lineages, the maximum ZOI observed between a pair of lineages was chosen as representative, which would mask the statistical signal of intra-lineage differences in antagonism. However, most cases of intra-lineage variation in sensitivity or antagonism (e.g. ZOI present for one isolate but not another in the same lineage) involved ZOIs near the limit of detection, and likely represented technical noise rather than true biological variation (Supplementary Figure 5). Thus, in order to look for mutations with functional effects on antagonism or sensitivity, we computed variance among all ZOI AUCs and considered only intra-lineage variations with >2 standard deviations between isolates in the same lineage. Still, in most cases, large intra-lineage variations were an artifact of background removal in image analysis (e.g. affected by adjacent spots or plate edges). For the four remaining cases (isolates 20.3, 37.3, 49.5, and 70.3), the possibility of parallel mutations was explored by examining 1) single nucleotide variants (SNVs) from Baker et al^6^, 2) genomic gains and losses from Baker et al^6^ and 3) small insertion-deletion and frameshift mutations identified in this study using Breseq^80^. Lineages 49 and 70 exhibited intra-lineage variation in sensitivity to lineage 34, with a single isolate out of five isolates showing sensitivity in each case (Supplementary Figure 3), but shared no similar mutations.

Lineages 20 and 37 exhibited intra-lineage variation in sensitivity to the same set of five antagonist lineages. Lineage 58 carried a similar sensitivity profile, though all tested isolates had the same sensitivity profile (Supplementary Figure 3). Parallel gene-disrupting mutations identified in vraFG pathway despite there being only 198 SNVs across lineage 20 and 128 SNVs across lineage 37. Lineage 58 was found on the same subject as isolate 20.3 carried a similar sensitivity phenotype and had SNVs in the vraG and dltABCDX pathways out of only 78 SNVs. Seeing parallel mutations in this pair of genes across all three lineages is unlikely to happen by chance (*P* < 0.0004, Binomial Test). Tree structures for these lineages indicated derived sensitivity rather than derived resistance (Supplementary Figure 16).

### Treemaking and distance calculations

Phylogenetic trees were generated in Baker et al^6^. In that study, Widevariant^6^ was used to tabulate core genome SNVs without signals of recombination. RaXML v8.2.12 was used to construct a species-wide maximum parsimony tree from a representative isolate for each lineage and DNApars v3.5c was used to construct a lineage-wide maximum parsimony tree from all isolates of selected lineages.

For computing pairwise distances, an average amino acid identity table was constructed from a representative single assembly from each lineage (for *S. epidermidis*) or species using EzAAI^81^. An average nucleotide identity table was constructed for all Staphylococcus isolates using FastANI^82^ from isolate assemblies.

### Statistical analysis of antagonism frequency and genomic similarity

We binned interactions by the pairwise amino acid identity or average nucleic acid identity of their corresponding pairs (Supplementary Figure 6) and computed a linear correlation between antagonism frequency or antagonism strength (AUC of ZOI) and genomic similarity. For comparing antagonism likelihood between categorical groupings, we shuffled species, phylogroup, or agr-type labels in permutation tests (Figure 2). Note that the observed differences would be significant under Fisher’s Exact test (e.g. for same vs different species, *P* = 0.001) but due to the structure in the interaction matrix, the independence assumption does not hold, and any differences are relatively small.

### MIC and growth assays

MIC and growth curve assays were performed on sub-cultured log-phase cells diluted to a starting OD of 0.0001. MIC assays were performed in 96-well plates (Greiner BioOne), in 200 µL TSB per well, shaking for 18 h overnight at 37°C. Growth curves were performed in 150 µL of TSB, grown in a LogPhase600 plate reader (Agilent) at 37°C, shaking at 800 RPM for 18h, sealed with Breathe-Easy plate seals (Research Products International 248738).

### Selection for reversion mutants

Idiosyncratic isolates 20.3 and 37.3 were grown overnight in TSB in 12 replicate 200 µL cultures. Each overnight culture was plated on TSA containing 64 µg/mL of polymyxin B and grown for 24 h at 37°C. The experiment was repeated a second time for isolate 20.3 using a single 25 mL culture grown in a flask and plating 2 mL per each of 12 plates on 150 mm petri dishes, but no resistant isolates were found. Resistant colonies 37.3-r1 and 37.3-r2 were grown in TSB 64 µg/mL polymyxin B, and the vraF and vraG sequences were confirmed by Sanger sequencing and whole genome sequencing. The phenotype for both 37.3-r1 and 37.3-r2 is the same, indicating that off-target mutations are unlikely to be responsible.

### Phage Isolation and Plaque Assay

A 5 mL subculture of *S. epidermidis* isolate 32.1 was grown shaking for 2 h at 37°C, then mitomycin C was added to a final concentration of 0.8 µg/mL. After 6 h of growth, when OD was observed to decrease, cells were pelleted at 2100 x g for 10 mins, and supernatant was filtered using a 0.22 µm filter. To determine plaquing counts, 100 µL diluted filtrates were mixed with 100 µL OD 0.5 subcultures of each *S. epidermidis*, then mixed with 3 mL of 0.5% agar and plated. We obtained plaques on lineage 37 isolates (Supplementary Figure 18), but did not obtain any plaques on isolates from lineage 20 or 58.

### Detection of phage in metagenomics

A 100 µL aliquot of the filtered supernatant from mitomycin-C treated culture of isolate 32.1 (containing the phage we refer to as v341) was used for DNA extraction, short-read sequencing, and assembly (see Sequencing and genome assembly above), resulting in a 42kb contig with 95% identity to published phage genomes targeting *S. epidermidis*^83^. Metagenomic reads generated in Baker et al^6^ were mapped against the v341 contig using Bowtie2. Reads from subjects 1PB (the source of lineage 32) and 1PA mapped to the v341 contig, as well as from subjects 7AA, 7AB, and 7PA, but not from subjects containing vraFG mutants (2PA, 2PB, and 5PA). As it is possible that these subjects may have had a similar phage in the past, we cannot conclude whether or not this specific phage or a relative thereof is responsible for any potential *in vivo* selection for vraFG mutations.

## Data availability

All sequencing data generated in previous work^6^ is available on the NCBI Sequence Read Archive under Bioproject # PRJNA1052084. All sequencing data generated in this work is available under Bioproject #PRJNA1215987. All images from the interaction screen are available on Figshare with doi: 10.6084/m9.figshare.25726482. All other data is included in the manuscript, supplemental tables, or code base (Code availability).

## Code availability

All code needed to perform the bioinformatic pipeline, image analysis pipeline, and reproduce the figures and analysis in this manuscript is available on GitHub at https://github.com/cpmancuso/Sepidermidis-antagonism.

**Supplementary Figure 1:**
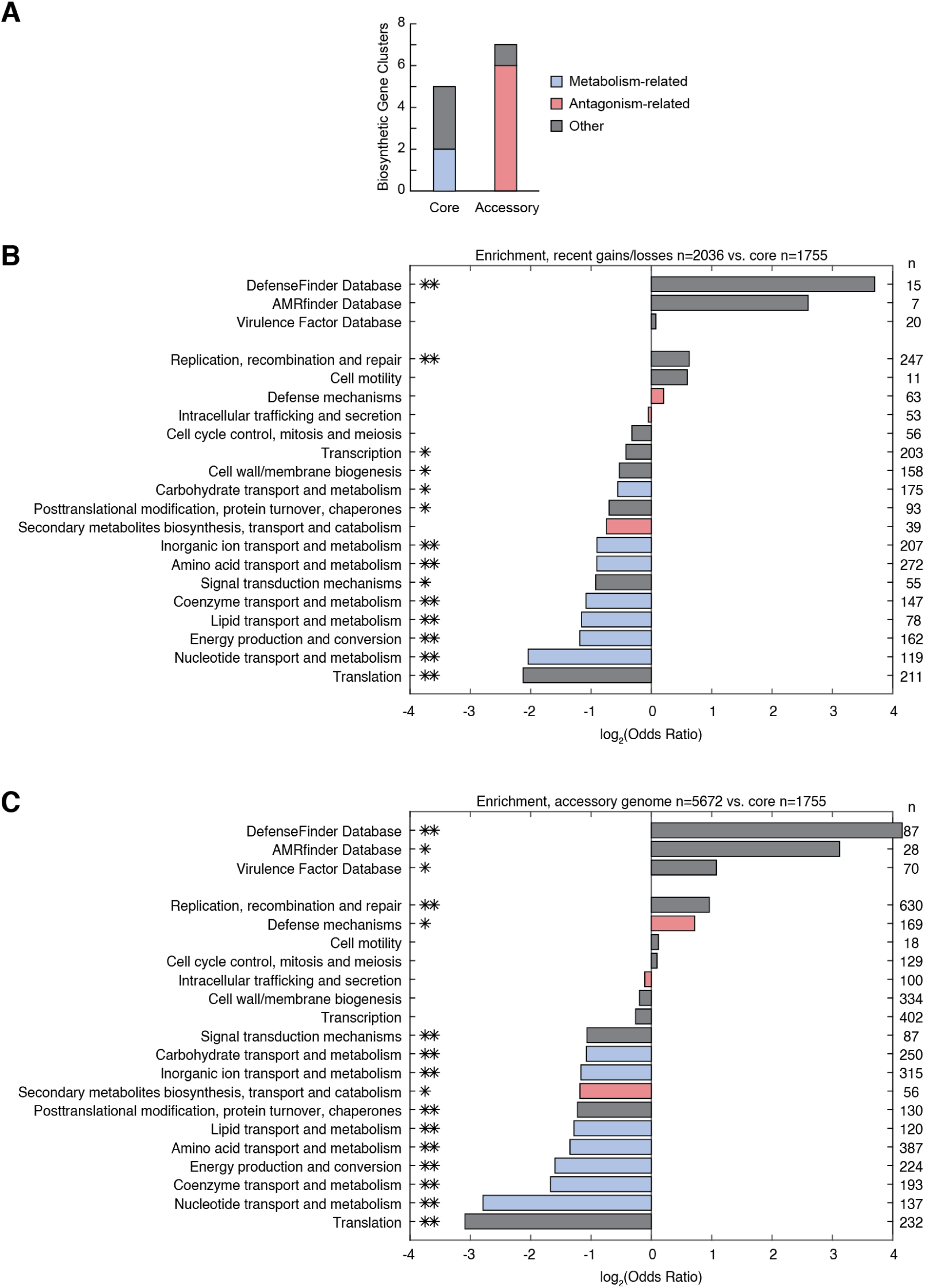
Antimicrobial BGCs and defense genes are enriched in accessory genomes. (A) We constructed a pangenome for *S. epidermidis* isolates in our collection and annotated genes and biosynthetic gene clusters (BGCs). Antagonism-related BGCs were more frequent in accessory genomes, whereas metabolism-related BGCs were found in the core genome. (B) We identified regions of the genome with copy number variation within a lineage, indicating recently gained or lost regions. We compared the relative enrichment of COG terms and hits from functional annotation databases among genes in the core genome and recent gain/loss regions. (C) As in B, but comparing genes in the core and accessory genomes. COG categories denoted in blue are metabolism associated, whereas antimicrobial production and resistance genes may be found in red categories. Significant enrichment is indicated by an asterisk (one for naive p-value < 0.05, two for Bonferroni corrected p-value < 0.05), and the number of genes annotated in each category is indicated to the right of the figure.

**Supplementary Figure 2:**
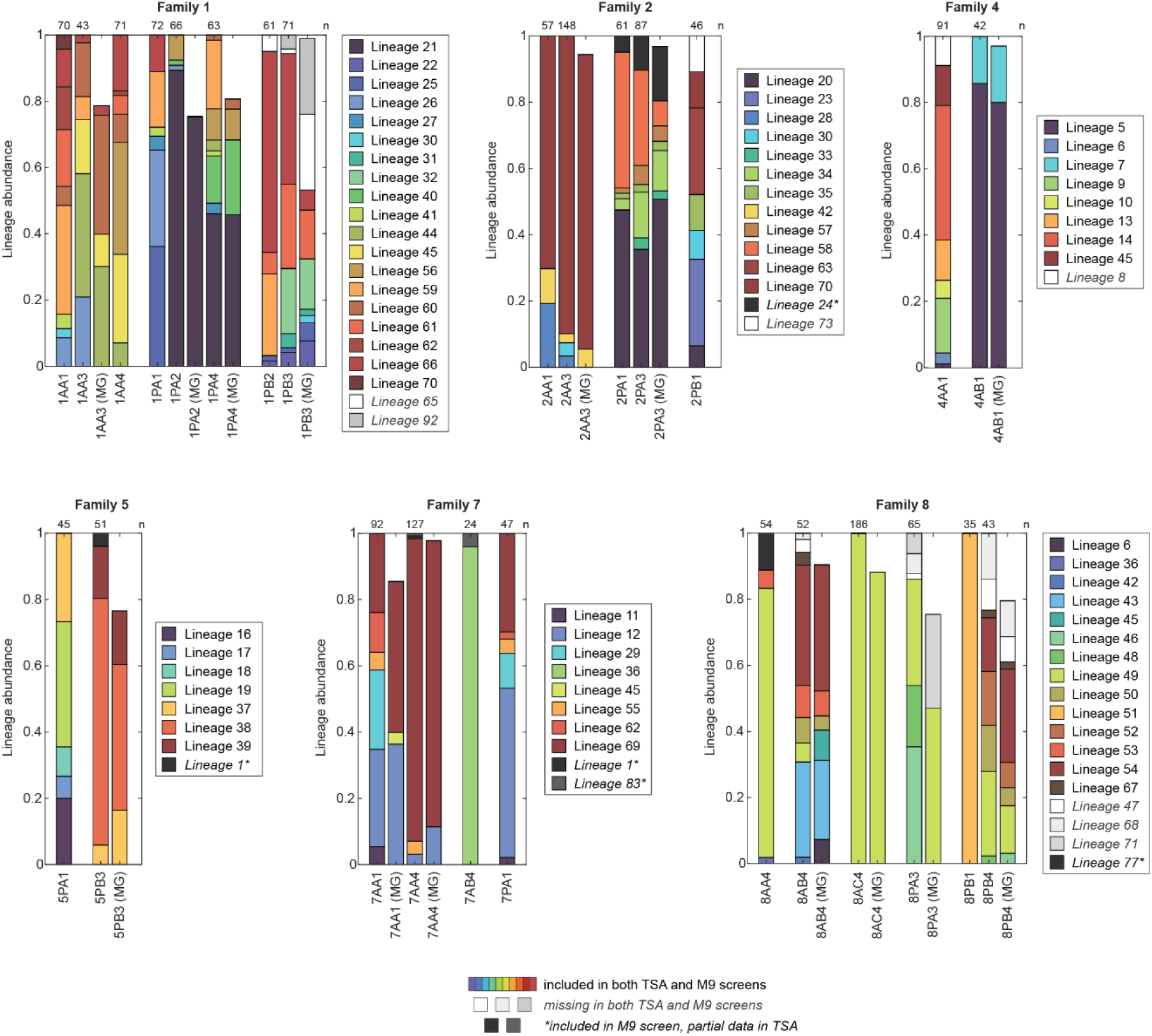
Representative isolates capture the *S. epidermidis* lineage diversity of individuals and families. In a previous study^6^, up to 96 isolates of *S. epidermidis* were cultured from each of 23 subjects at each timepoint sampled during a 1.5-year time period. Six families were selected for inclusion in this study. Relative lineage abundance, as determined from isolates, is plotted for each sample, organized by subject and family. PHLAME^69^ was applied to metagenomic samples from the same subject (denoted as MG), which confirmed that lineage abundances estimated by culturing did not have significant culture bias^6^; note that uncalled portions of metagenomic samples (space above bars) can reflect uncertainty in PHLAME due low coverage rather than the true absence of lineages. We used isolate-based lineage abundances because deep metagenomic coverage of *S. epidermidis* was not available for every sample. Sample labels indicate the family, subject, and sampling timepoint (e.g. 1AA3 indicates Family 1, subject 1AA, timepoint 3). The number of *S. epidermidis* colonies isolated from each person is indicated above each bar. Lineages represented in the TSA antagonism screen in this study are indicated in color, while unrepresented lineages are indicated in greyscale. Some additional lineages were included in a follow-up screen M9+S screen (see Methods); these are indicated by darker grey colors (denoted by *). Colors do not carry over across families, lineages are generally restricted to a family. Across all samples, the average proportion of lineage composition represented by isolates included in the TSA antagonism screen was 96%.

**Supplementary Figure 3:**
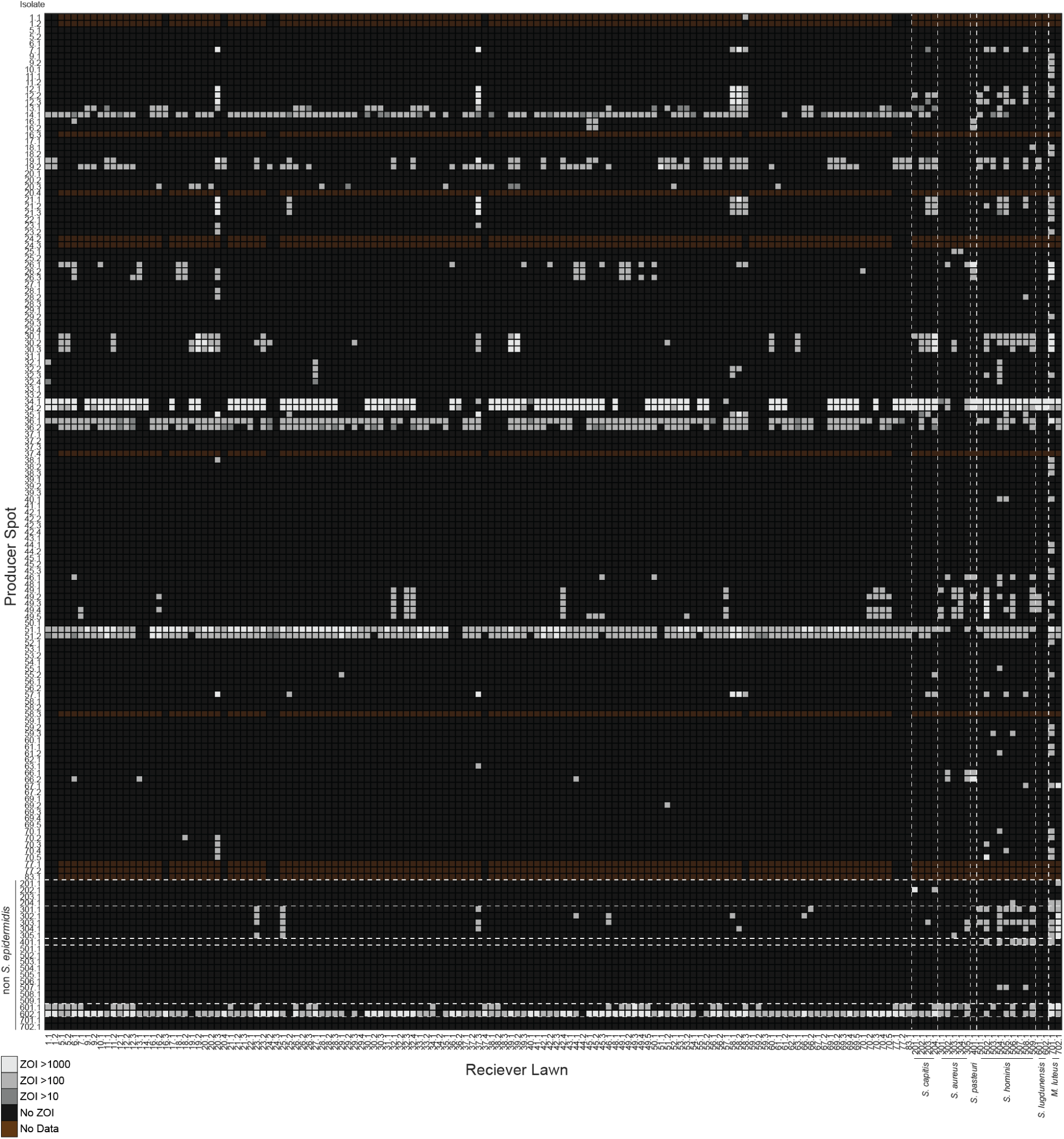
Antagonism and sensitivity generally cluster at the lineage level. We measured 21,025 pairwise interactions between 145 isolates that passed inclusion criteria (15 replicate cultures were also included, totaling 25,600 interactions). The heatmap depicts the AUC of the ZOI calculated for pairwise interactions between *S. epidermidis* and other skin microbiome isolates in TSA. Each row and column indicate an isolate (in contrast to Figure 2A where rows indicate lineages). Multiple isolates from a lineage are denoted by decimals (e.g. 20.1, 20.2, 20.3). Isolates are sorted by phylogenetic similarity, with dashed lines to indicate species boundaries. Each isolate of species other than *S. epidermidis* (labels > 200, see x-axis for details) was treated as a separate lineage. Interactions were shared among all isolates of the same lineage in ∼97% of cases, so *S. epidermidis* isolates were dereplicated into lineages for other analyses unless otherwise noted. Nine isolates were added after the preliminary TSA screen, and thus are lacking some data points, indicated by brown squares. Unless otherwise noted, these isolates with partial data were excluded from statistical analysis.

**Supplementary Figure 4:**
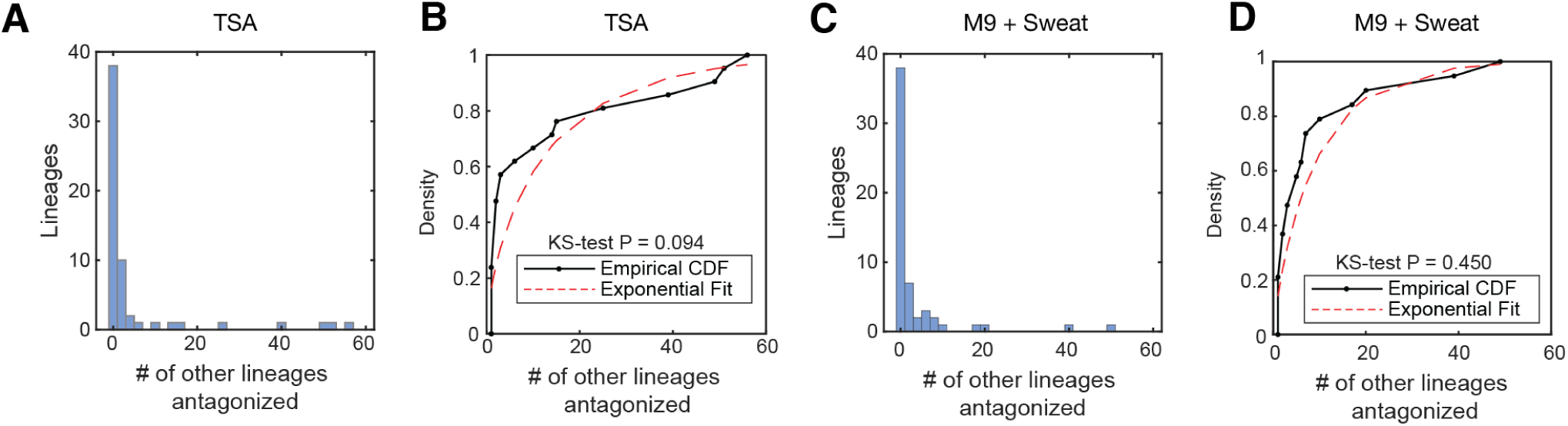
Antagonism proportion is exponentially distributed. (A) Histogram of showing the distribution of the number of other lineages antagonized by a given lineage. (B) The distribution of antagonism proportion is not distinguishable from an exponential fit, where p-value represents the result of a Kolmogorov– Smirnov test to compare the empirical CDF to the exponential fit. (C) As in A, but for M9 + Sweat screen. (D) as in B, but for M9 + Sweat screen. As antagonism proportion fits an exponential distribution, we did not exclude any lineages as outliers.

**Supplementary Figure 5:**
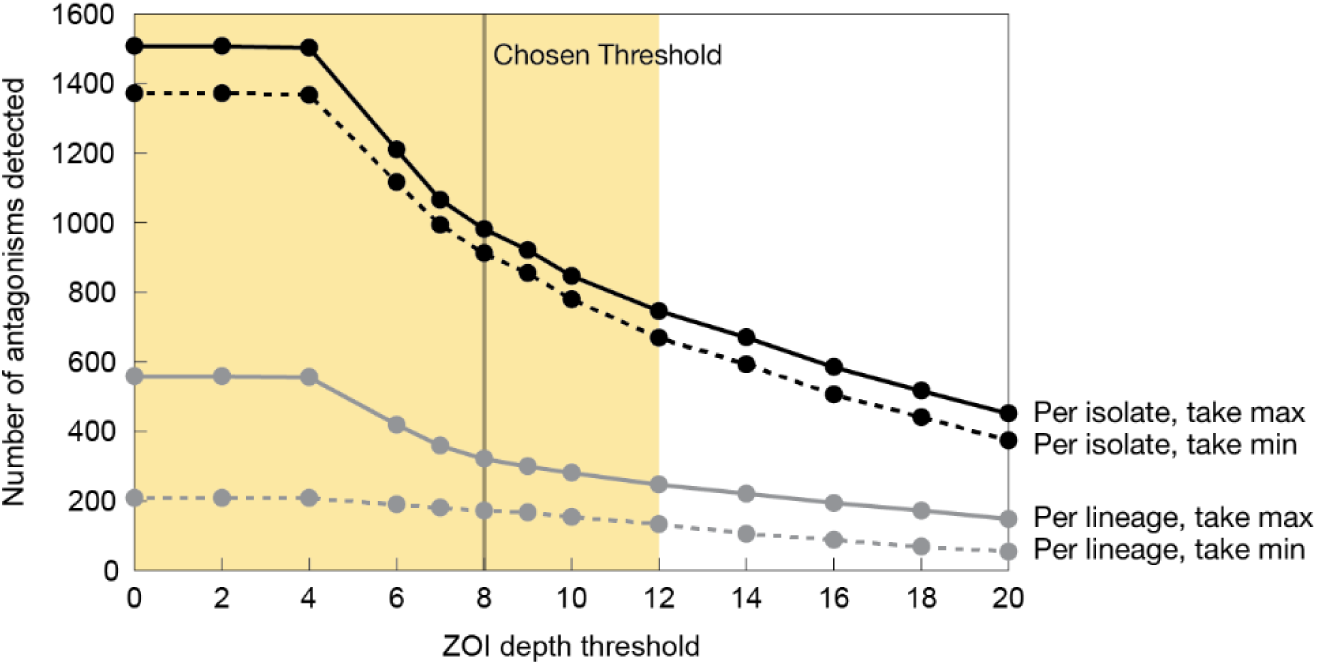
Results are robust to alternative choice of thresholds. The threshold cutoffs were chosen to maximize agreement between ZOI calls made by eye vs. by our image analysis pipeline; here we varied the threshold based on ZOI depth, the maximum change in pixel intensity at any radial point surrounding the culture spot. At low and moderate thresholds (yellow background), the major results of our study remain significant. However, at more permissive thresholds, the number of ZOIs increases exponentially, potentially indicating spurious ZOI calls. At higher thresholds, legitimate cases of antagonism begin to be excluded and mean antagonism frequency drops low enough that some statistical tests (especially Supplementary Figure 11E) begin to return non-significant results. Separately, the chosen ZOI Depth threshold is approximately the optimum value to minimize the percent difference between two possible lineage dereplication methods (maximizing vs minimizing the number of antagonisms). These findings suggest that our results are not confounded by noisy detection of antagonism or dependent solely on antagonisms of a certain strength.

**Supplementary Figure 6:**
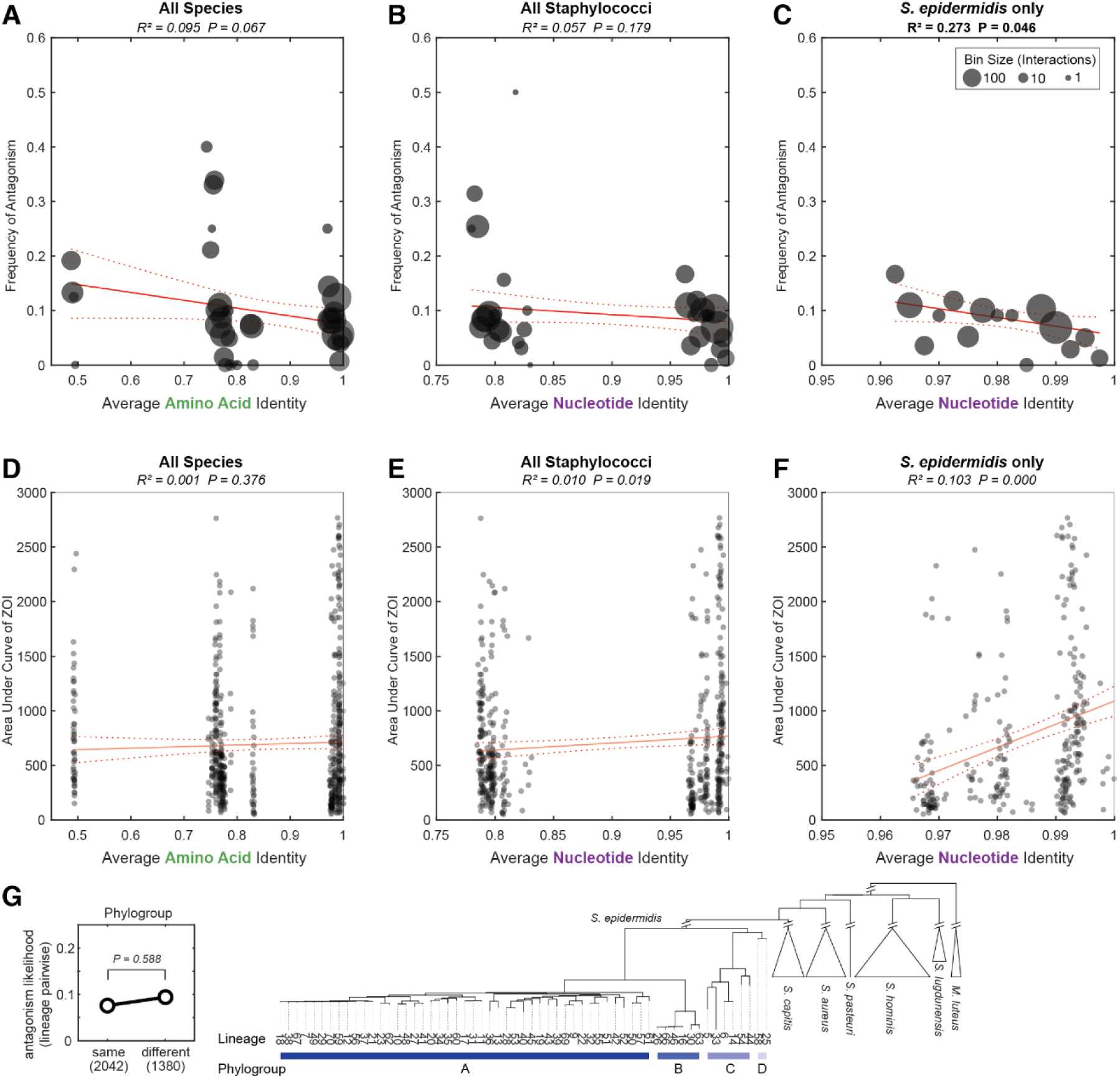
Antagonism frequency and strength show limited variation at different phylogenetic distances. Rather than depicting genomic relatedness using a tree (Figure 2A), we calculated pairwise distances by different sequence similarities to understand whether antagonism and relatedness are correlated. (A) We binned interactions between pairs of lineages with similar average amino acid identities, then plotted the frequency of antagonism among pairs in each bin. (B) As in A, but binned according to average nucleotide identity. (C) As in B, but only for lineages of *S. epidermidis*, not other species. Antagonism frequency does not significantly vary with genomic distance in our collection, as both R^2^ and *P* are weak. (D) We plotted the strength of antagonism (AUC of ZOI) for each antagonistic interaction (excluding non-antagonistic interactions) against the average amino acid identity of the pair. (E) As in D, but against the average nucleotide identity. (F) As in E, but only for lineages of *S. epidermidis*, not other species. There is a significant but small increase in antagonism strength at high nucleotide identities among *S. epidermidis*. (G) The frequency of antagonism does not significantly differ between members of the same or different *S. epidermidis* phylogroup, according to a two-sided permutation test with shuffled group labels. The phylogeny at right depicts *S. epidermidis* phylogroup boundaries. Taken together, these results indicate that antagonism occurs at all levels of genomic relatedness, even between closely related lineages.

**Supplementary Figure 7:**
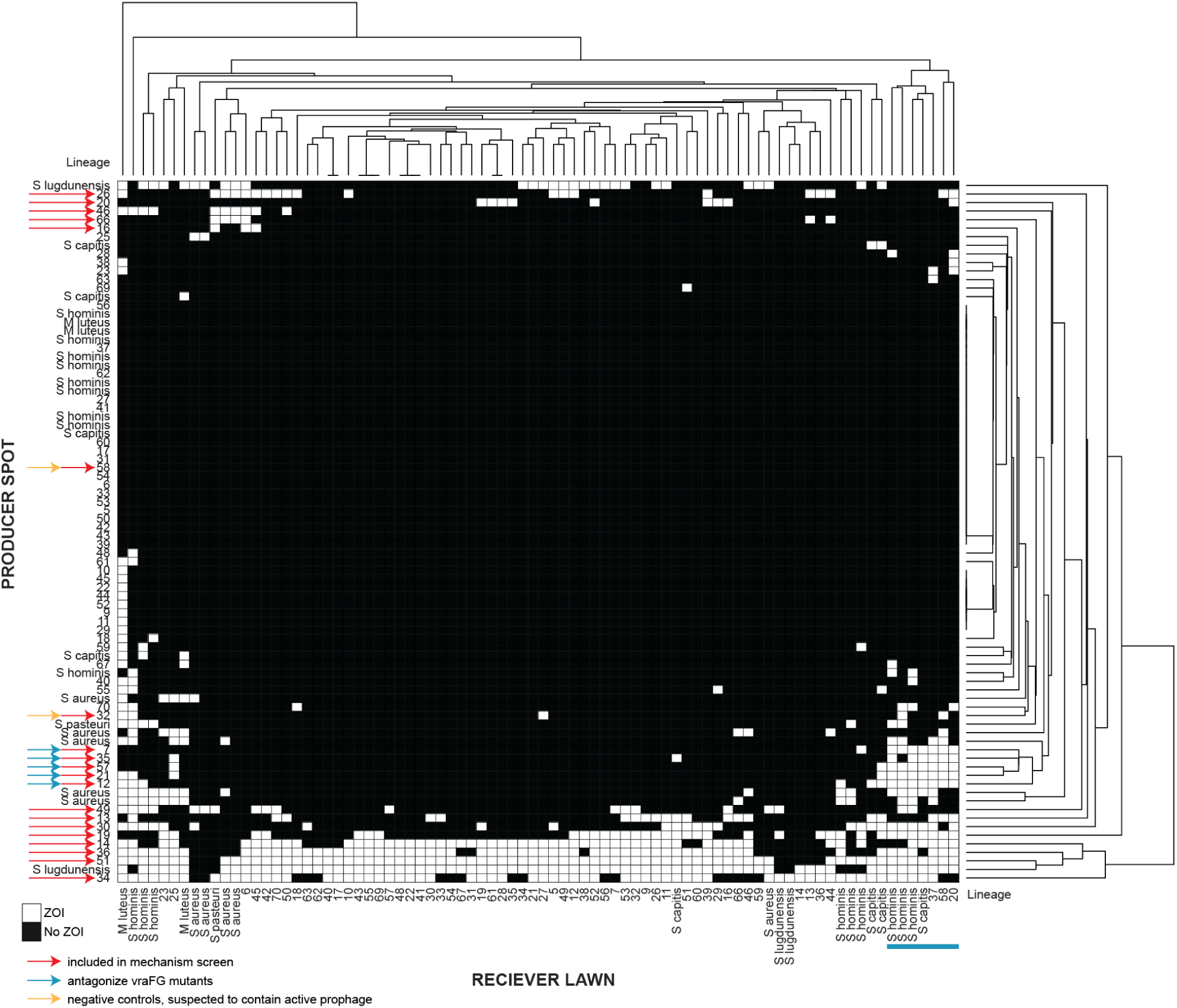
Wide variation in intra– and inter-species antagonism patterns. In order to check for shared mechanisms, the binary antagonism matrix was clustered by constructing dendrograms for antagonism and sensitivity (Methods). White boxes indicate antagonism by the producer spot against the receiver lawn. Red arrows indicate the lineages in our collection chosen for our mechanism screen (Supplementary Figure 8), which include 18 lineages with high antagonism frequencies plus 2 negative controls (which lacked antimicrobial production genes but contained suspected prophage, denoted by yellow arrows). We chose representative receiver bait lineages using the dendrogram of sensitivity to ensure that all interaction patterns were captured. Note that for this analysis, unlike other analyses in this study (Methods), antagonisms from idiosyncratic isolates that exhibit intra-lineage variation were not excluded. Lineages denoted by cyan arrows did not inhibit many *S. epidermidis*, with the exception of idiosyncratic lineages 20, 37, and 58, which cluster with some *S. hominis* and *S. capitis* isolates, denoted by a cyan line. It is possible that the antimicrobials behind these interactions are intended to target non-epidermidis species.

**Supplementary Figure 8:**
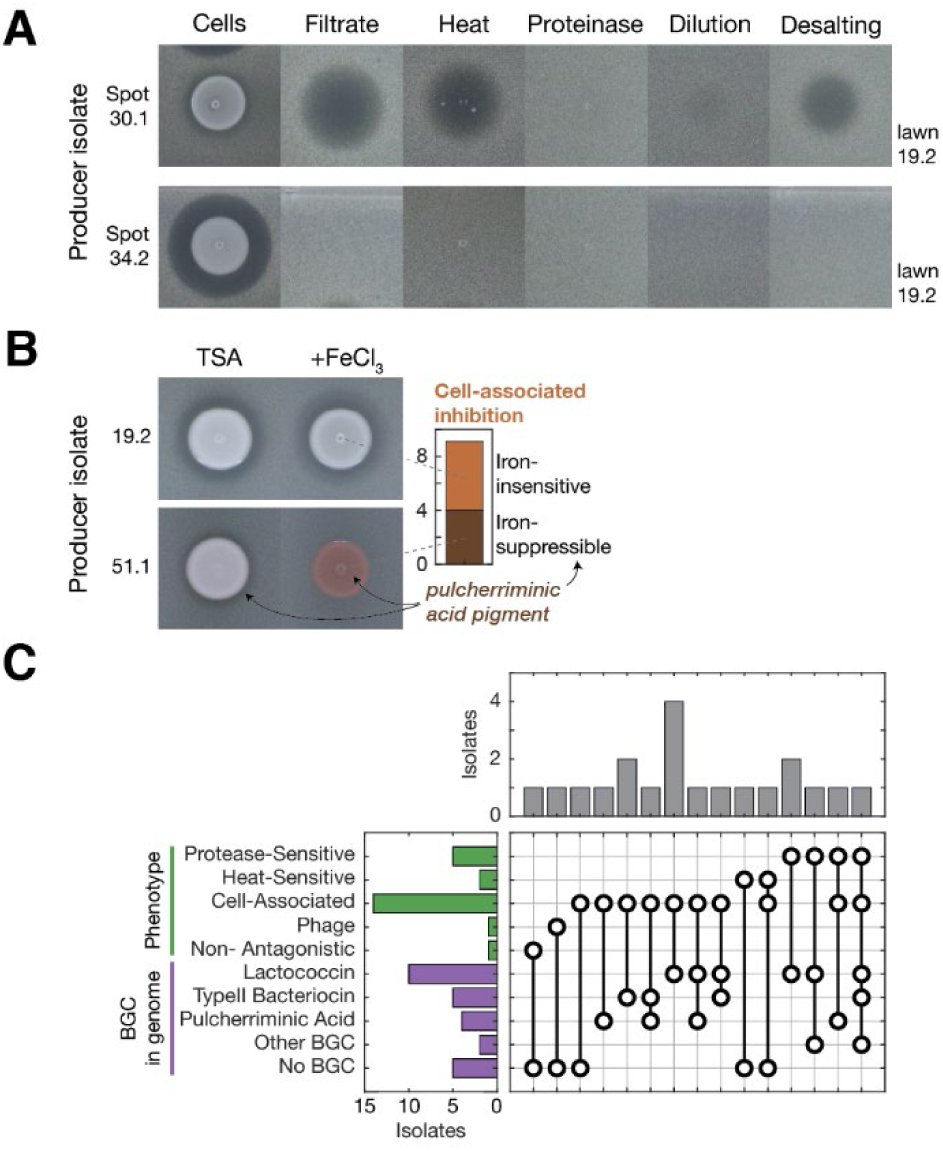
Warfare between *S. epidermidis* strains is mediated by diverse and unannotated effectors. (A) To explore the diversity of antagonism mechanisms, we screened cells and filtered supernatants from 18 antagonistic *S. epidermidis* isolates against a representative array of bait lawns chosen based on interaction profiles (as well as 2 negative controls; Supplementary Figure 7). Filtrates treated with heat or proteinase to degrade proteins and peptides, desalting to remove small molecules, or 1:10 dilution were also tested. Filtrates exhibited varied patterns, including antagonisms likely mediated by small peptide effectors (isolate 30.1) and some cell-associated antagonisms induced by neighbors (isolate 34.2) or both. (B) Some cell-associated antagonisms are iron-mediated, as indicated by the elimination of the interaction in media supplemented with excess FeCl_3_ (see isolate 51.1). All tested isolates displaying iron-suppressible antagonisms cells contained a biosynthetic gene cluster (BGC) for a product similar to pulcherriminic acid, an iron-sequestering pigment molecule which sabotages the iron supply^42^. (C) We annotated putative bacteriocin BGCs and found BGCs for known compounds (including gallidermin-like^37^, epicidin-like^38^, and lactococcin-like BGCs) as well as unknown compounds. However, we did not find a clear correspondence between BGC presence and the phenotype classified in our mechanism screen, potentially due to multiple inducible effectors being encoded in the genome, incomplete expression of BGCs in different media or environments, or targets not represented in our screen (e.g. *Cutibacterium acnes*).

**Supplementary Figure 9:**
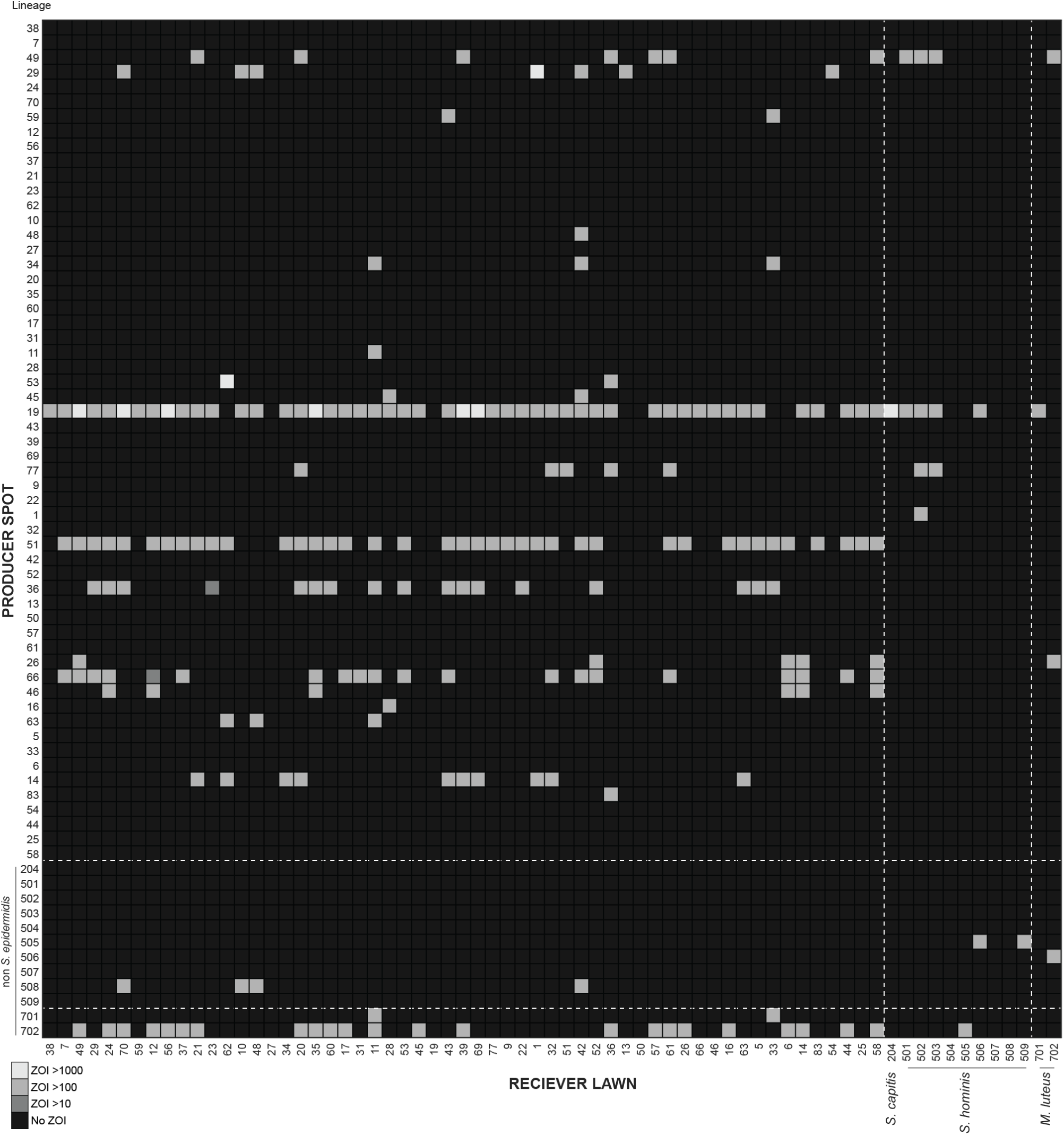
Antagonism and sensitivity at the lineage level measured in M9 with Artificial Sweat. We measured pairwise interactions between 114 isolates that passed inclusion criteria in M9 + Sweat media, then dereplicated *S. epidermidis* interactions to the lineage level (see Methods). The heatmap depicts the AUC of the ZOI calculated for pairwise interactions between *S. epidermidis* and other skin microbiome isolates. Each row and column indicate a lineage. Each isolate from species other than *S. epidermidis* (labels > 200, see x-axis for details) was treated as a separate lineage. See Supplementary Figure 10 for a comparison between media conditions.

**Supplementary Figure 10:**
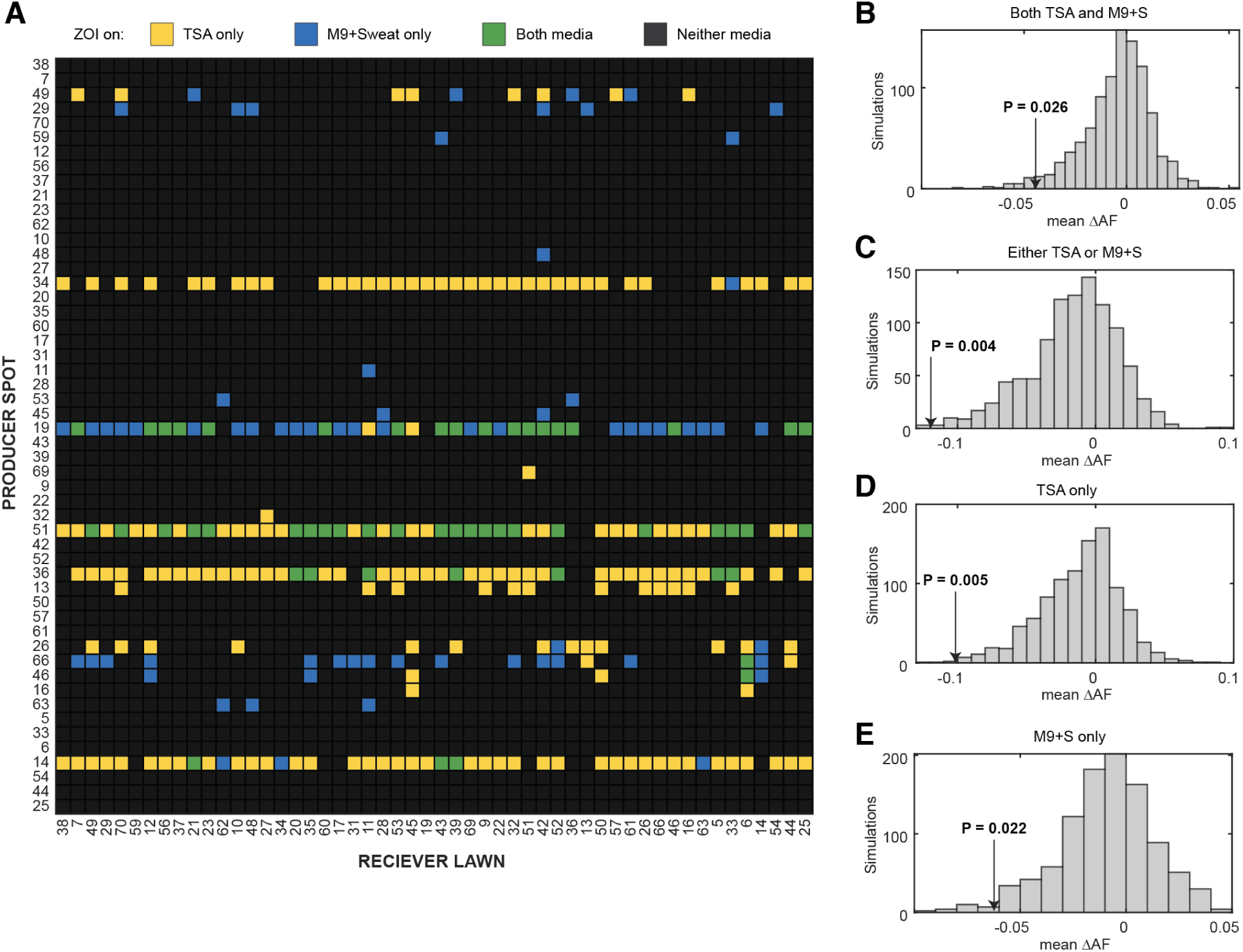
Antagonism differences between TSA and M9+S do not affect depletion of on-person antagonism. (A) We overlaid interactions from TSA onto the *S. epidermidis* interaction matrix generated in M9+S media, using color to indicate which media the antagonism appeared in. In total, 91% of interactions are the same in both media conditions, though antagonisms did vary between the two media, such that 23% of antagonisms observed in TSA were also observed in M9+S, and 61% of antagonisms observed in M9+S were observed in TSA. (B-D) Despite these differences, we observed a significant depletion of on-person antagonism (ΔAF, see Figure 3C, Methods) in permutation tests that compare our observed data compared to simulated data with shuffled lineage compositions on each subject.

**Supplementary Figure 11:**
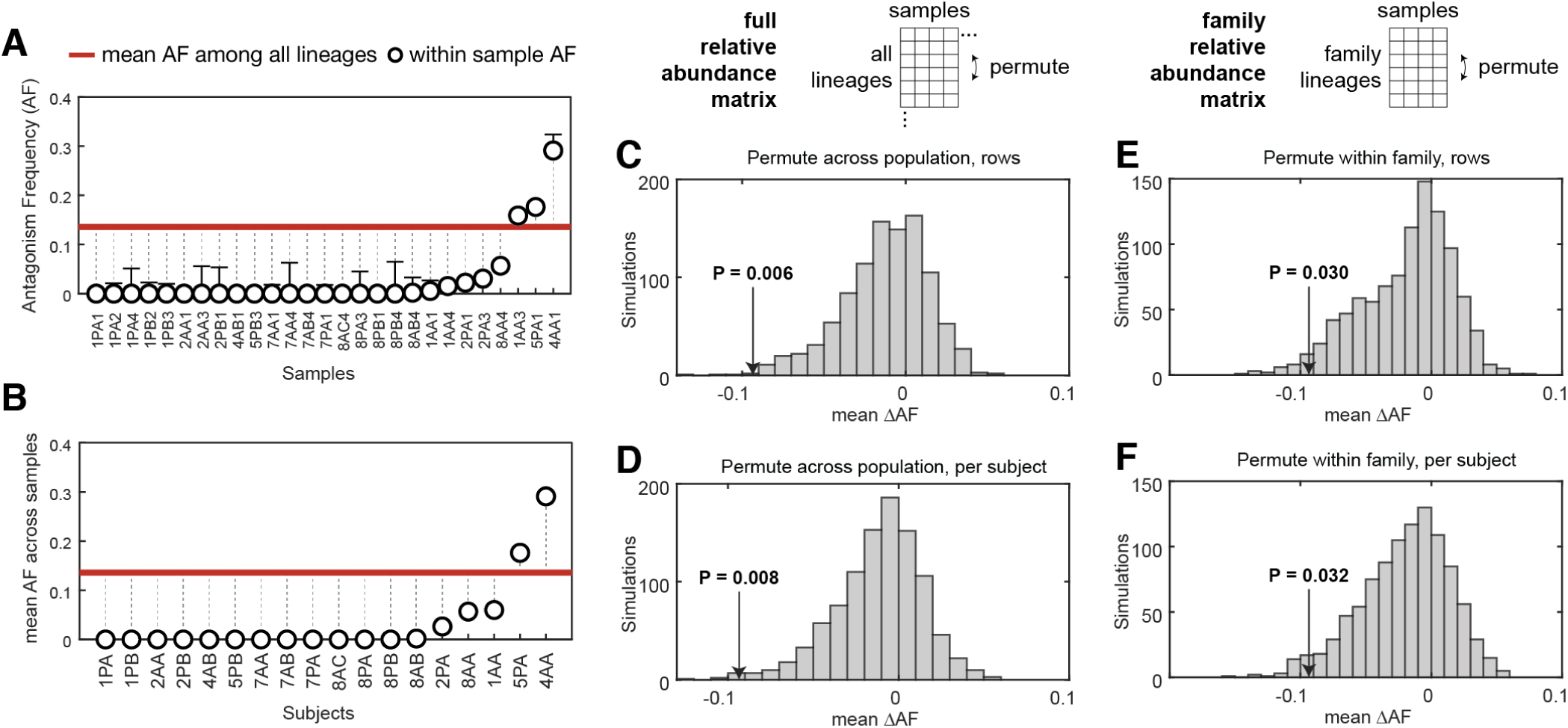
Antagonistic interactions are depleted in samples, individuals, and families. (A) Antagonism frequencies (AF) between co-resident lineages in samples appears depleted relative to the frequency across all *S. epidermidis* lineages (red). When lineages were present in a sample but were absent from our screen, we calculated the range of AF that would be expected based on the average rate of antagonism between any two isolates, depicted as error bars. This is likely an overestimate of the range, since on-person antagonism frequencies are lower than population-wide antagonism frequency. (B) We took the mean AF across samples of the same subject, and still found that mean within sample AF appeared to be depleted relative to the expected AF. Reproduced from Figure 3A. (C-F) Since t-tests are invalid for non-independent samples, we performed permutation testing by simulating scenarios that retained the structure in the interaction matrix and the composition of samples (e.g. uneven lineage distribution). For each simulation, we shuffled lineage labels in the relative abundance matrix and the p-value represents the fraction of simulations with less on-person antagonism than observed (black arrow indicates observed value of ΔAF). (C) The rows of the composition matrix were shuffled once across all lineages in our collection. This simulation tests for depletion of antagonism while maintaining the same degree of lineage sharing seen between individuals in the same family. Reproduced from Figure 3C. (D) The rows of the composition matrix were shuffled, AF values were calculated for samples of the first subject, then rows were shuffled again before each new subject. These simulations broke the relatedness between individuals in the same family, while retaining similarity between samples from the same subject, thereby testing depletion of antagonism under the assumption that lineages are acquired from the environment independent of family. (E) To test for depletion of antagonism under the assumption that one can only acquire strains from a family member, shuffling was only performed on submatrices composed only of lineages on a family were shuffled, by row, as in C. (F) As in F, but without maintenance of lineage sharing between family members; submatrices shuffled but by once per subject. Note that less shuffled conditions (E,F) will definitionally have higher p-values because of fewer possible permutations. Under all tested assumptions, antagonism is depleted among strains present on the same person at the same time.

**Supplementary Figure 12:**
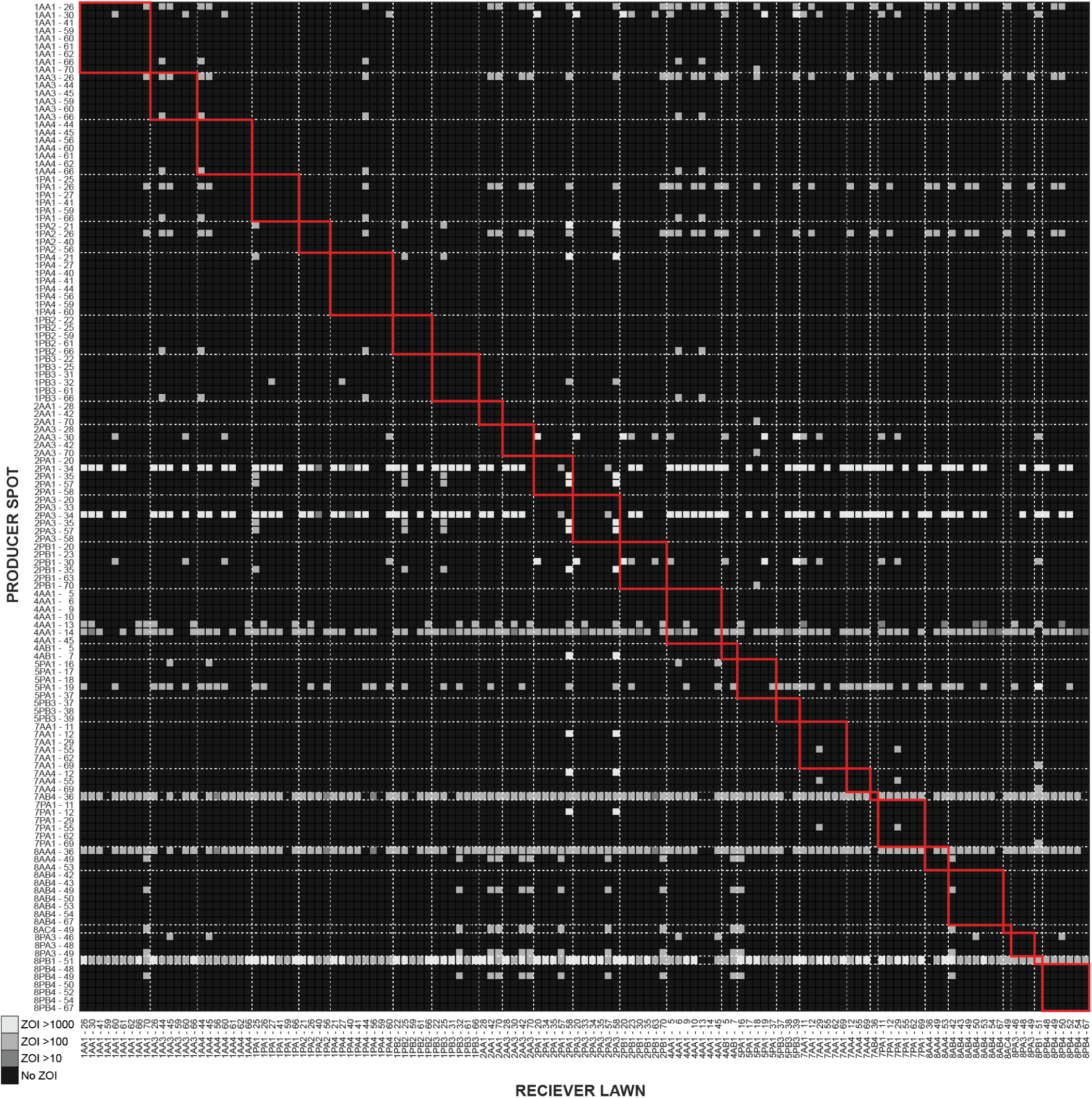
Subjects colonized by antagonists have different lineage compositions than family members. The heatmap depicts the AUC of the ZOI observed on TSA for pairwise interactions between *S. epidermidis* lineages found on each subject. Each row and column indicate a lineage present in a sample, labelled by the sample in which the lineage was found (e.g. 1AA1 – 26 indicates lineage 26 on subject 1AA timepoint 1. Note that whenever lineages were found at multiple timepoints or on multiple subjects, the data for that lineage is repeated (e.g. 1AA1-26 and 1PA1-26 are the same data). Dashed lines indicate the boundaries between samples. Red boxes denote areas of the interactions matrix that represent interactions between co-resident lineages. A subset of this data is shown in Figure 3D, highlighting two families with subjects colonized by antagonist lineages.

**Supplementary Figure 13:**
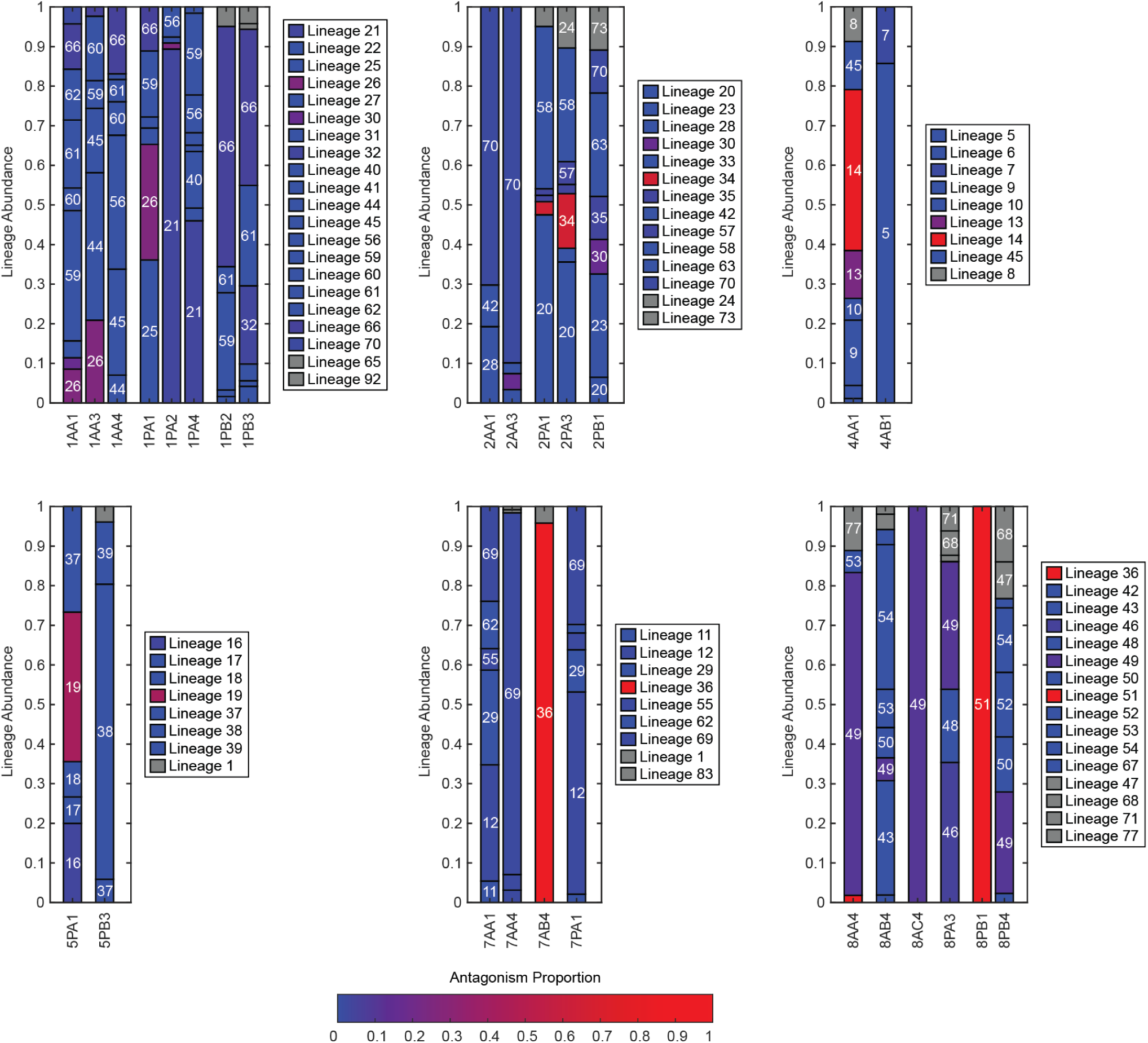
Subjects colonized by antagonists have different lineage compositions than family members. Relative abundance plots where each lineage is colored by the proportion of other lineages it antagonizes on TSA, with blues representing non-antagonistic lineages and reds representing highly antagonistic lineages. Lineages are labelled by number in white text and are plotted in the same order as in Supplementary Figure 2. Grey lineages have missing data, and therefore an antagonism proportion cannot be calculated. Note that antagonism proportion is relative to other lineages across the study, not just within a family. Some families contain few antagonistic lineages (e.g. family 1), whereas most families contain at least one highly antagonistic lineage.

**Supplementary Figure 14:**
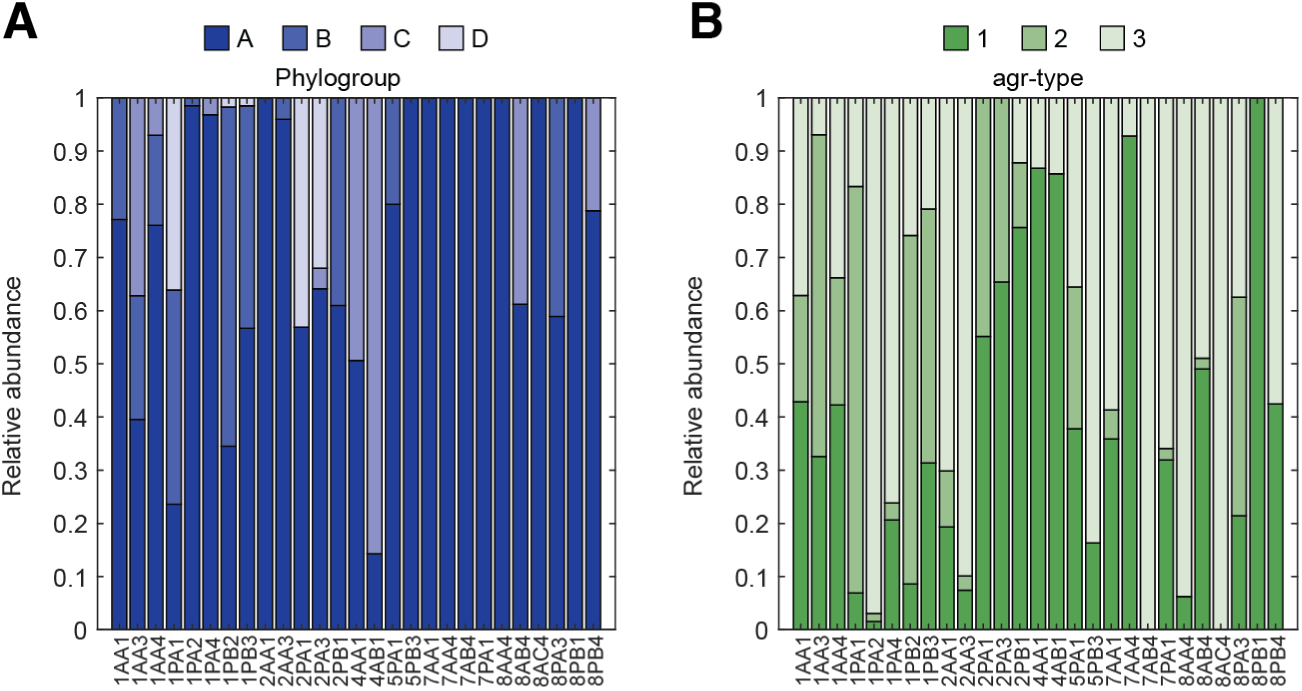
Multiple phylogroups and quorum sensing types are coresident on facial skin. (A) Lineages were grouped into phylogroups according to phylogenetic structure as described previously^6^, see Supplementary Figure 6G. On many subjects, multiple phylogroups coexist on the face at the same time. (B) The agrD sequence was annotated for each lineage on each subject and classified into quorum sensing types. Again, we found that on many subjects, multiple agr types coexist on the face at the same time. Co-residence between different phylogroups and between incompatible quorum sensing types suggest that there is not competitive exclusion on the basis of phylogenic dissimilarity or quorum sensing.

**Supplementary Figure 15:**
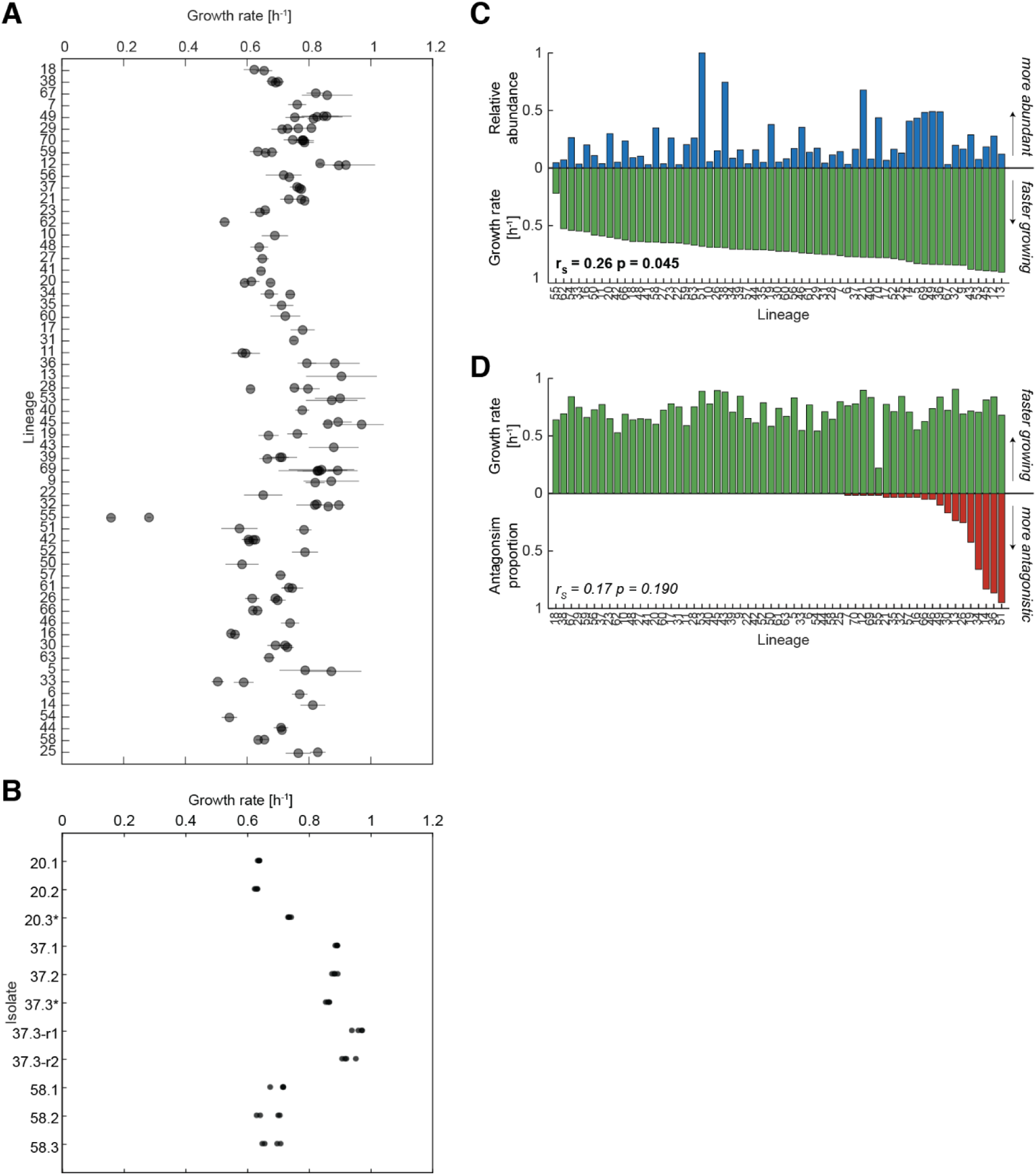
Growth rate and fitness variation among *S. epidermidis*. (A) We measured growth rate in liquid culture in TSB for all isolates in our collection (Methods). We calculated the median growth rate for each isolate in a lineage. Bars denote standard deviation (from technical duplicates of biological triplicates). (B) The growth rate for each isolate of idiosyncratic lineages 20, 37, and 58 (technical duplicates of biological quadruplicates) is shown. Growth rate varies between vraFG mutants (indicated with *) and wild-type isolates of each lineage, but without a directional trend. Reversion mutants of 37 have faster growth rates than 37.3, leaving the basis of the vraFG mutations unresolved. (C) Lineages that grow faster than other lineages *in vitro* (green) reach higher abundance on individuals (blue), Spearman’s rank correlation. Note that growth *in vivo* may vary significantly due to differences between nutrient composition of skin and TSB. (D) Lineages that antagonize a higher fraction of other lineages (red) do not grow faster or slower than other lineages (green), Spearman’s rank correlation.

**Supplementary Figure 16:**
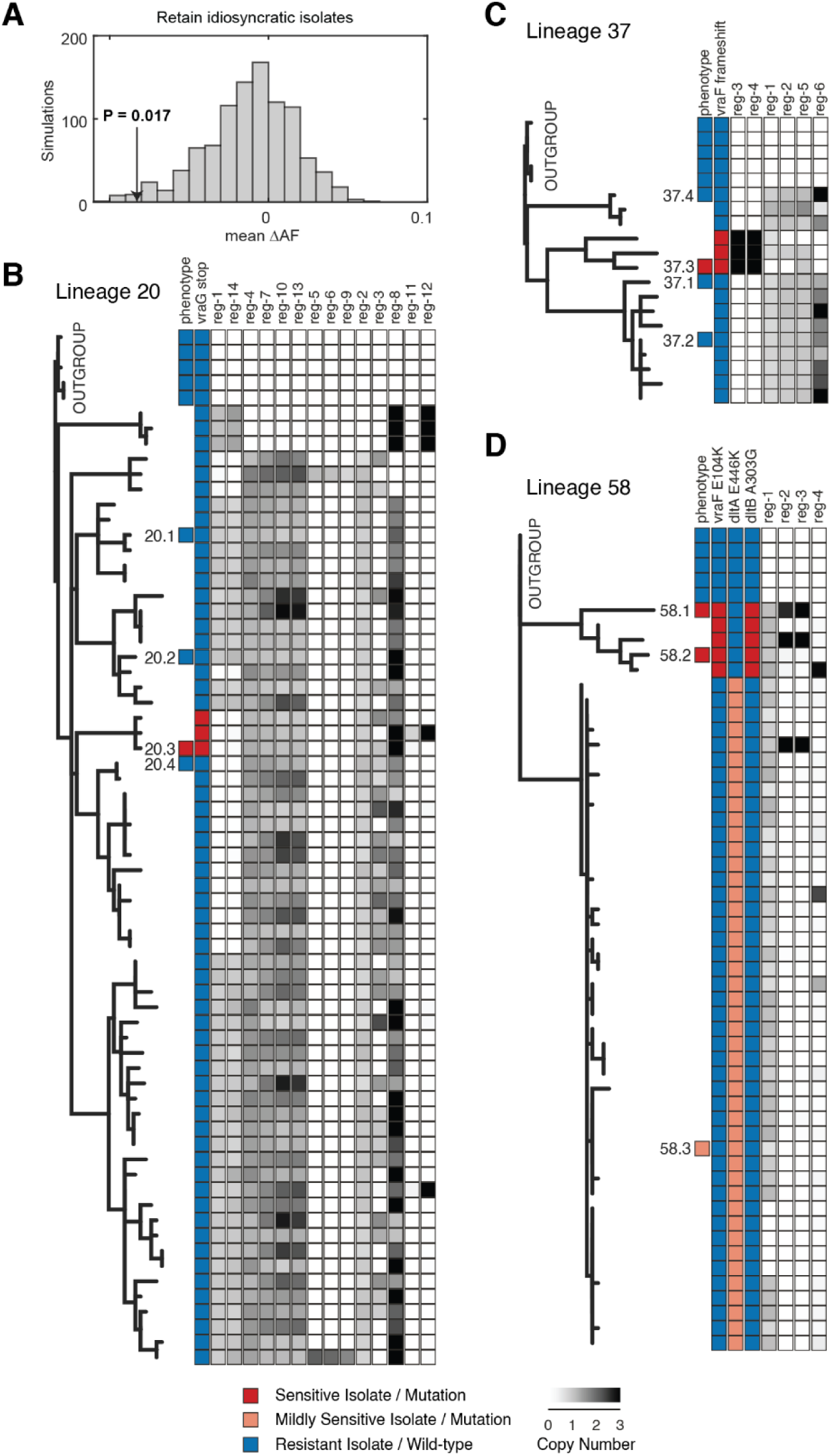
Phylogenies show rare cases of derived sensitivity to antimicrobials. (A) Four “Idiosyncratic” isolates (20.3, 37.3, 49.5, and 70.3) exhibited large differences in antimicrobial sensitivity compared to other members of their respective lineages. Since each represented a minority of their lineages, these isolates were excluded from other analyses presented in this study. However, on-person antagonisms remain depleted even if these isolates are retained (using simulations as in Fig. 3C, TSB). (B) Two idiosyncratic isolates shared a sensitivity pattern with lineage 58 (Figure 4A, Supplemental Figure 7), prompting a search for parallel mutations. We plotted the sensitivity phenotype against whole genome single nucleotide polymorphism trees constructed for each lineage. Each isolate on the tree was annotated with any *vraFG* and *dlt* operon mutations and/or genomic region gains or losses (colored by copy number relative to core genome). The sensitive phenotype (red/orange) correlated with *vraF* and *vraG* mutations in (B) lineage 20, (C) lineage 37, and (D) lineage 58. Presence or absence of genomic regions have no apparent trend in regards to the idiosyncratic sensitivity phenotype. While the *vraFG* mutants form a small fraction of their respective lineages, these isolates were found at multiple timepoints over six months apart, indicating that they are not a transient part of the population.

**Supplementary Figure 17:**
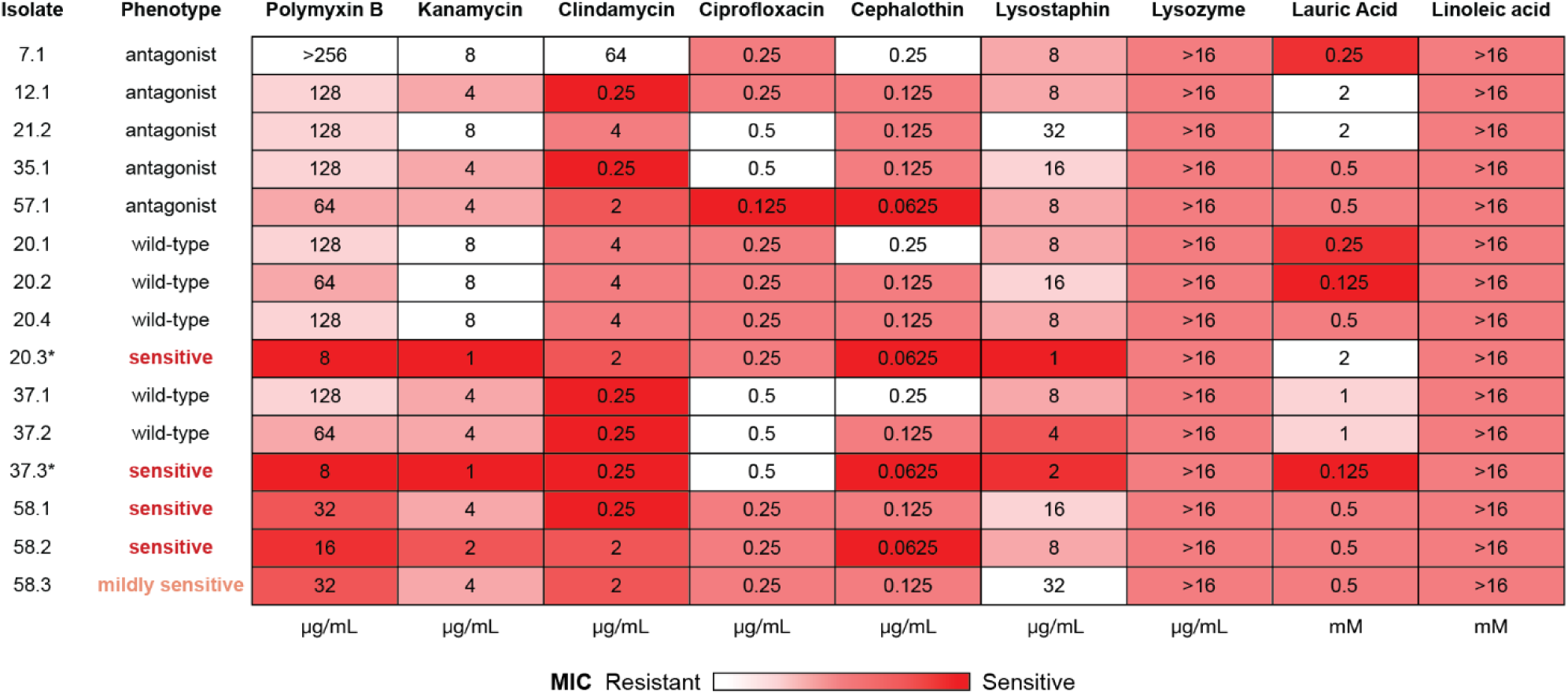
Mutations cause correlated sensitivity to cationic antimicrobials and lysostaphin but not to other stresses. Heatmap depicting the Minimum Inhibitory Concentration for antibiotics, lysis enzymes, and antimicrobial fatty acids. Polymyxin B (+5) and kanamycin (+4) are more positively charged than clindamycin (+1), ciprofloxacin (0), or cephalothin (–1). Idiosyncratic isolates 20.3 and 37.3 are more sensitive to cationic antimicrobials and lysostaphin than other isolates in their lineages. Both *vraFG* and *dlt* operon genotypes in lineage 58 (see Supplementary Figure 16) share a similar MIC pattern to these idiosyncratic isolates, especially in response to Polymyxin B.

**Supplementary Figure 18:**
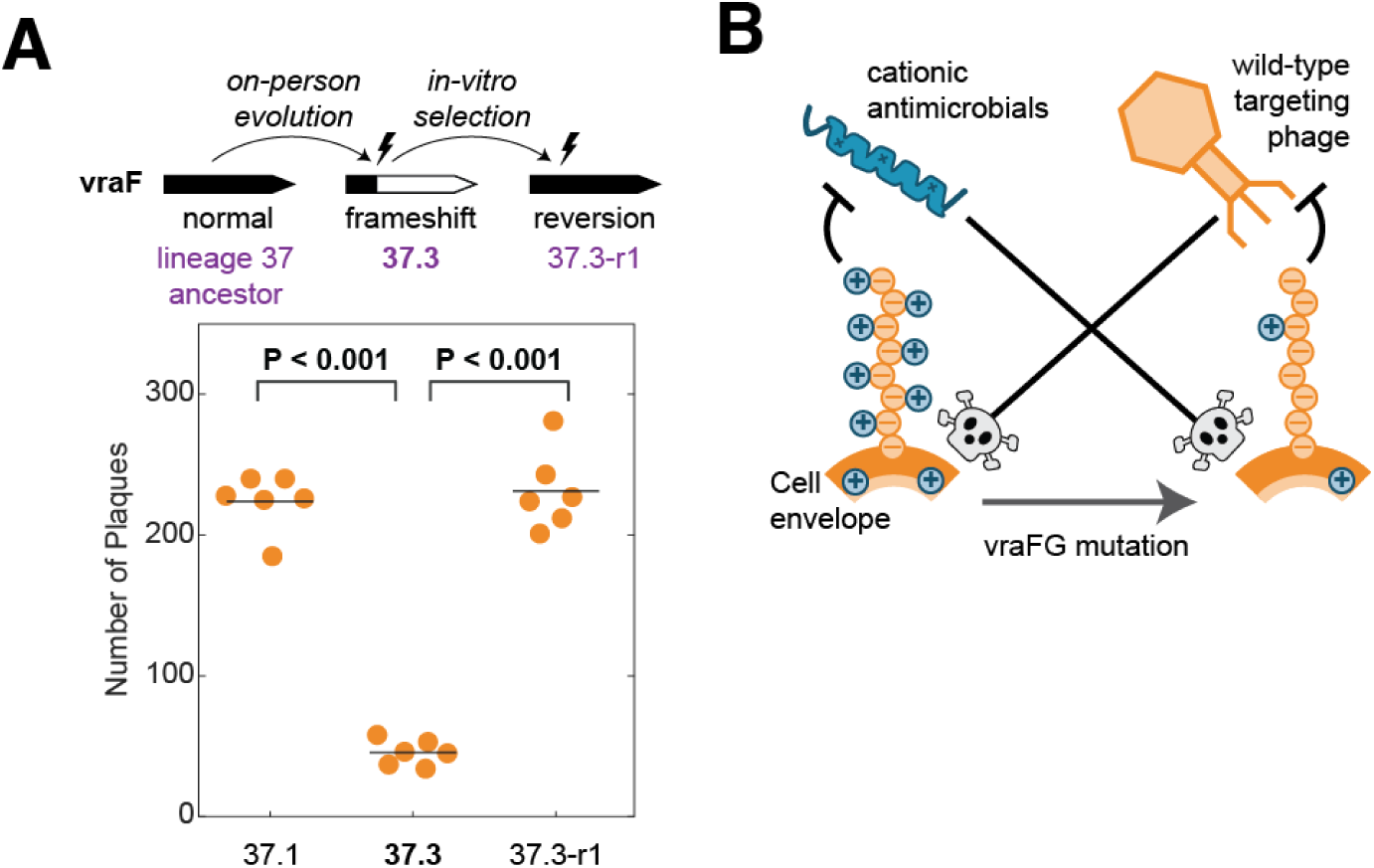
Mutation in vraF yields partial phage resistance in potential tradeoff. (A) Despite other sensitivities (Figure 4; Supplementary Figure 17), the vraF-frameshift isolate (37.3) was less susceptible to phage infection than isolates with full-length vraF. However, we did not find this phage on subjects carrying lineage 37 (Methods). (B) Nevertheless, together with prior results from literature^49,52^, these results are consistent with a conceptual model in which the vraFG complex could reduce cell envelope modifications, weakening resistance to cationic antimicrobials but providing protection against some phage.

## Acknowledgements

We thank the participants of the original study and C. Dickinson, S. Friedhoff, M. Hirsch, S. Zuckermann, and J. Schuler for assistance with study recruitment. We thank J. Zhou, Z. Zhang, S. Zhang, and S. Levine for advice on experimental design. We thank all members of the Lieberman Lab for advice on this project and feedback on the manuscript.

This work was funded by The James H. Ferry, Jr. Fund for Innovation in Research Education, Colgate-Palmolive Company, and NIH grant 1DP2GM140922 (all to T.D.L).

## Author Information

Contributions: C.P.M. and T.D.L. conceived of the project. J.S.B. collected and sequenced the samples. C.P.M., A.D.T., and T.D.L. designed the experiments. C.P.M. carried out the experiments and conducted image processing and statistical analyses. C.P.M., J.S.B., E.Q., A.D.T., and I.O.B. conducted bioinformatic analyses. T.D.L. secured funding for the project. The original manuscript was drafted by C.P.M. and T.D.L.; all authors reviewed and edited the final manuscript.

